# Loss of the extracellular matrix molecule tenascin-C leads to absence of reactive gliosis and promotes anti-inflammatory cytokine expression in an autoimmune glaucoma mouse model

**DOI:** 10.1101/2020.04.28.064758

**Authors:** Susanne Wiemann, Jacqueline Reinhard, Sabrina Reinehr, Zülal Cibir, Stephanie C. Joachim, Andreas Faissner

## Abstract

Previous studies demonstrated that retinal damage correlates with a massive remodeling of extracellular matrix (ECM) molecules and reactive gliosis. However, the functional significance of the ECM in retinal neurodegeneration is still unknown. In the present study, we used an intraocular pressure (IOP) independent experimental autoimmune glaucoma (EAG) mouse model to examine the role of the ECM glycoprotein tenascin-C (Tnc).

Wild type (WT ONA) and Tnc knockout (KO ONA) mice were immunized with an optic nerve antigen (ONA) homogenate and control groups (CO) obtained sodium chloride (WT CO, KO CO). IOP was measured weekly and electroretinographies were recorded at the end of the study. 10 weeks after immunization, we analyzed retinal ganglion cells (RGCs), glial cells and the expression of different cytokines in retina and optic nerve tissue in all four groups.

IOP and retinal function was comparable in all groups. Although less severe in KO ONA, WT and KO mice displayed a significant loss of RGCs after immunization. Compared to KO ONA, a significant reduction of βIII-tubulin stained axons and oligodendrocyte markers was noted in the optic nerve of WT ONA. In retinal and optic nerve slices, we found an enhanced GFAP^+^ staining area of astrocytes in immunized WT. In retinal flat-mounts, a significantly higher number of Iba1^+^ microglia was found in WT ONA, while a lower number of Iba1^+^ cells was observed in KO ONA. Furthermore, an increased expression of the glial markers *Gfap, Iba1, Nos2* and *Cd68* was detected in retinal and optic nerve tissue of WT ONA, whereas comparable levels were observed in KO ONA post immunization. In addition, pro-inflammatory *Tnfa* expression was upregulated in WT ONA, but downregulated in KO ONA. Vice versa, a significantly increased anti-inflammatory *Tgfb* expression was measured in KO ONA animals.

Collectively, this study revealed that Tnc plays an important role in glial and inflammatory response during retinal neurodegeneration. Our results provide evidence that Tnc is involved in glaucomatous damage by regulating retinal glial activation and cytokine release. Thus, this transgenic EAG mouse model offers for the first time the possibility to investigate IOP-independent glaucomatous damage in direct relation to ECM remodeling.

## 1 Introduction

Glaucomatous neurodegeneration is characterized by a progressive loss of retinal ganglion cells (RGCs) and their axons, which form the optic nerve. The molecular mechanisms of RGC degeneration are not fully understood. In addition to increased intraocular pressure (IOP), immunological processes, glial activation, and remodeling of extracellular matrix (ECM) constituents are associated with glaucoma. In regard to the immune system, studies also indicate an alteration in serum antibodies against various retinal proteins in glaucoma patients with a normal IOP (Tezel et al., 1998; Wax et al., 2001; Wax, 2011). The connection between an immune response and glaucoma disease with the characteristic loss of RGCs was already demonstrated in an experimental autoimmune glaucoma (EAG) rat model. Here, glaucomatous damage was induced by immunization with ocular proteins (Laspas et al., 2011; Joachim et al., 2013). Furthermore, a pathological upregulation of specific ECM components could be demonstrated in this model (Reinehr et al., 2016). However, the relationship between a change in ECM components and glaucoma pathogenesis is still unknown.

The ECM consists of several molecules, including proteoglycans and glycoproteins, and controls cellular key events such as adhesion, differentiation, migration, proliferation as well as survival (Hynes, 2009; Theocharidis et al., 2014; Faissner and Reinhard, 2015; Krishnaswamy et al., 2019; Roll and Faissner, 2019; Theocharis et al., 2019). ECM molecules can provide an inhibitory environment for neural regeneration and migration in the retina (Reinhard et al., 2015). A dramatic remodeling of ECM constituents has already been described after ischemia and glaucomatous damage (Reinhard et al., 2017a; Reinhard et al., 2017b). For instance, several studies reported a dysregulation of the glycoprotein tenascin-C (Tnc) during neurodegeneration. An upregulation of Tnc has been described in a glaucoma animal model (Johnson et al., 2007) and in patients with open-angle glaucoma (Pena et al., 1999). Tnc is also a key regulator of the immune system and plays an important role during neuroinflammation and glial response (Jakovcevski et al., 2013; Dzyubenko et al., 2018; Wiemann et al., 2019). In the retina, this glycoprotein is specifically expressed by amacrine and horizontal cells (D’Alessandri et al., 1995). Moreover, expression of Tnc by astrocytes is regulated via cytokines secreted by microglia (Smith and Hale, 1997; Haage et al., 2019). Microglia play an important role during neurodegenerative and neuroinflammatory processes (Glass et al., 2010). Their activation is characterized by an enhanced proliferation, migration, phagocytosis, and increased expression levels of neuroinflammatory molecules (Langmann, 2007; Wolf et al., 2017). The neurotoxic M1-subtype has an amoeboid morphology and releases pro-inflammatory signaling molecules, like tumor necrosis factor-alpha (TNF-α) and inducible nitric oxide synthase (iNOS) (Harms et al., 2012; Varnum and Ikezu, 2012; Silverman and Wong, 2018). In contrast, the M2-phenotype is characterized by a morphology with ramified processes and the expression of anti-inflammatory cytokines such as the transforming growth factor-beta (TGF-β) (De Simone et al., 2004; Colton, 2009; Ramirez et al., 2017).

In this study, we used a Tnc deficient EAG mouse model to further analyze the importance of Tnc during retinal neurodegeneration and neuroinflammatory outcomes in glaucoma disease. We immunized wild type (WT) and Tnc knockout (KO) mice with an optic nerve antigen homogenate (ONA) and examined retinal and optic nerve damage as well as macro- and microglial activity. Furthermore, we determined the expression pattern of pro- and anti-inflammatory cytokines. The present study was undertaken to address the role of Tnc in glaucomatous damage, retinal glial activation, myelination and inflammatory cytokine release.

## 2 Materials and Methods

### 2.1 Animals

Animals were housed under a 12 h light-dark cycle and had free access to chow and water. All procedures were approved by the animal care committee of North Rhine-Westphalia, Germany and performed according to the ARVO statement for the use of animals in ophthalmic and vision research. For the experiments, male and female 129/Sv WT and *Tnc* KO mice (Forsberg et al., 1996) were used at 6 weeks of age.

### 2.2 Immunization

WT (WT ONA) and KO (KO ONA) mice were immunized with ONA (1 mg/ml) mixed with incomplete Freund’s adjuvants (FA) and 1 µg pertussis toxin (PTx; both Sigma Aldrich, St. Louis, MO, USA) according to the previously described pilot study (Reinehr et al., 2019). FA acted as an immunostimulatory and PTx was given to ensure the permeability of the blood retina barrier. PTx-application was repeated 2 days after immunization. Booster injections containing half of the initial dose were given 4 and 8 weeks after initial immunization. The control groups (WT CO; KO CO) were injected with 1 ml sodium chloride (B. Braun Melsungen AG, Melsungen, Germany), FA and PTx. 10 weeks after immunization, retinae and optic nerves were explanted for immunohistochemistry, quantitative real time PCR (RT-qPCR), and Western blot analyses. For RT-qPCR and Western blot, we pooled retinal and optic nerve tissue of both eyes.

### 2.3 Intraocular pressure measurements

IOP measurements were performed before immunization in WT and KO mice at 5 weeks of age with a rebound tonometer (TonoLab; Icare; Oy; Finland; n = 16/group) as previously described (Schmid et al., 2014; Reinhard et al., 2019). After immunization, IOP was measured weekly in all groups until the end of the study. Before IOP measurement mice were anesthetized with a ketamine/xylazine mixture (120/16 mg/kg). Both eyes were analyzed, and 10 readings of each eye were averaged (n = 8/group).

### 2.4 Electroretinogram recordings

Scotopic full-field flash electroretinograms (ERG) recordings (HMsERG system, OcuScience, Henderson, NV, USA) were taken 10 weeks after immunization in all groups (n = 5/group) as previously described (Reinhard et al., 2019). Mice were dark-adapted and anaesthetized with a ketamine/xylazine mixture (120/16 mg/kg). Scotopic flash series with flash intensities at 0.1, 0.3, 1.0, 3.0, 10.0, and 25.0 cd/m^2^ were recorded. Electrical potentials were analyzed with the ERGView 4.380R software (OcuScience) using a 150 Hz filter before evaluating a- and b-wave amplitudes.

### 2.5 Immunohistochemistry and confocal laser scanning microscopy

Eyes and optic nerves were dissected and fixed in paraformaldehyde (PFA) for 1 day, dehydrated in sucrose (30 %), and embedded in Tissue-Tek freezing medium (Thermo Fisher Scientific, Cheshire, UK). Retinal cross-sections and optic nerve longitudinal sections (16 µm) were cut with a cryostat (CM3050 S, Leica) and transferred onto Superfrost plus object slides (Menzel-Glaeser, Braunschweig, Germany). First, slices were blocked with 1 % bovine serum albumin (BSA; Sigma-Aldrich), 3 % goat serum (Dianova, Hamburg, Germany), and 0.5 % Triton™-X-100 (Sigma-Aldrich) in phosphate-buffered saline (PBS) for 1 hour (h) at room temperature (RT). Afterwards, the primary antibodies were diluted in blocking solution and incubated overnight at RT (Table 1). Sections were washed 3 times in PBS and incubated for 2 h with adequate secondary antibody (Dianova, Hamburg, Germany; Table 1) solution without Triton™-X-100. Cell nuclei were detected with TO-PRO-3 (1:400; Thermo Fisher Scientific). The retinal and optic nerve slices were analyzed with a confocal laser-scanning microscope (LSM 510 META; Zeiss, Göttingen, Germany). 2 sections per slide, 4 images per retina (400x magnification) and 3 images per optic nerve (200x magnification) were captured (n = 4-5/group). In addition, a 630x magnification was used for colocalization staining in optic nerve sections with antibodies against CC1 (coiled-coil 1) and Olig2 (oligodendrocyte transcription factor 2). Accordingly, 4 images were taken per slide (n = 5/group).

**Table 1:** List of primary and secondary antibodies to examine RGCs, macro- and microglial cell types in retinae and optic nerves via immunohistochemistry. Primary and secondary antibodies, including species, clonality/type, dilution and source/stock number/RRID were shown.

| Primary Antibody | Species, Clonality/Type | Dilution | Source (Stock no.)/RRID | Secondary Antibody | Species, Type | Dilution/Source |
| --- | --- | --- | --- | --- | --- | --- |
| Brn-3a<br>(Flat-mount/<br>Retina) | Goat,<br>polyclonal,<br>IgG | 1:300 | Santa Cruz<br>(Sc-31984)<br>RRID:AB_2167511 | Anti-goat<br>Cy3 | Donkey,<br>IgG | 1:250/<br>Dianova |
| CC1<br>(Optic nerve) | Mouse,<br>monoclonal,<br>IgG | 1:100 | Abcam<br>(Ab16794)<br>RRID:AB_443473 | Anti-mouse<br>Cy2 | Goat,<br>IgG | 1:250/<br>Dianova |
| GFAP<br>(Retina/Optic<br>nerve) | Mouse,<br>monoclonal,<br>IgG | 1:300 | Sigma-Aldrich<br>(G3893)<br>RRID:AB_477010 | Anti-mouse<br>Cy2 | Goat,<br>IgG | 1:250/<br>Dianova |
| Iba1<br>(Flat-mount) | Rabbit,<br>polyclonal, IgG | 1:250 | WAKO<br>(019-19741)<br>RRID:AB_839504 | Anti-rabbit<br>Cy2 | Donkey,<br>IgG | 1:250/<br>Dianova |
| MBP<br>(Optic nerve) | Rat,<br>monoclonal,<br>IgG | 1:250 | Bio-Rad<br>(MCA409)<br>RRID:AB_325004 | Anti-rat<br>Cy2 | Goat,<br>IgG | 1:250/<br>Dianova |
| Olig2<br>(Optic nerve) | Rabbit,<br>polyclonal,<br>IgG | 1:400 | Merck<br>(AB9610)<br>RRID:AB_570666 | Anti-rabbit<br>Cy3 | Goat,<br>IgG | 1:250/<br>Dianova |
| βIII-tubulin<br>(Optic nerve) | Mouse,<br>monoclonal,<br>IgG2b | 1:300 | Sigma-Aldrich<br>(T 8660)<br>RRID:AB_477590 | Anti-mouse<br>Cy2 | Goat,<br>IgG | 1:250/<br>Dianova |

Laser lines and emission filters were adjusted using the Zeiss ZEN black software. Cropping of the images was done using Coral Paint Shop Pro X8 (Coral Corporation, CA, USA). Masked evaluation of the staining signal was performed with ImageJ software (ImageJ 1.51w, National Institutes of Health; Bethesda, MD, USA) as previously described (Reinehr et al., 2016; Reinehr et al., 2018). Images were converted into grey scales and the background was subtracted. Then, the lower and upper threshold values was determined for each image (Table 2). The percentage of the area fraction was measured using an ImageJ macro as previously described (Reinehr et al., 2018). This analysis was performed for immunohistochemical stainings against βIII-tubulin, glial fibrillary acidic protein (GFAP) and myelin basic protein (MBP). Cell countings were done for immunopositive Brn3a^+^ cells in retinal cross-sections and for Olig2^+^/CC1^+^ cells in optic nerve slices. Values were transferred to Statistica software and the WT CO group was set to 100 % (V13.3; StatSoft (Europe), Hamburg, Germany).

**Table 2:** Adjustments for the ImageJ macro. Data of background subtractions and upper/lower thresholds were present to determine the immunoreactive area [%].

| Protein | Tissue | Background Subtraction | Lower Threshold | Upper Threshold |
| --- | --- | --- | --- | --- |
| βIII-tubulin | Optic nerve | 50 | 29.45 | 82.18 |
| GFAP | Retina | 20 | 25.0 | 76.54 |
|  | Optic nerve | 50 | 27.50 | 78.0 |
| MBP | Optic nerve | 50 | 7.62 | 62.45 |

### 2.6 Quantification of RGCs und microglia in retinal flat-mounts

Eyes were enucleated and fixed in 4 % PFA for 1 h at 4°C. The retinae were dissected from the eye and prepared as flat-mounts (n = 9/group). The tissue was fixed again in 4 % PFA for 5 minutes and washed 3 times in PBS. Flat-mounts were blocked in 1 % BSA, 3 % donkey serum and 2 % Triton™-X-100 in PBS for 1 h at RT. Next, incubation was performed with the RGC specific marker Brn3a (brain-specific homeobox/POU domain protein 3a) (Xiang et al., 1996; Nadal-Nicolas et al., 2009) and microglia marker Iba1 (ionized calcium-binding adapter molecule 1) (Ito et al., 1998) for 2 days at 4°C. Following PBS washing (3 × 20 minutes), flat-mounts were incubated with secondary antibodies donkey anti-goat Cy3, donkey anti-rabbit Alexa Fluor 488 and TO-PRO-3 (1:400) in blocking solution without Triton™-X-100 for 2 h at RT. Microscopic images were captured using Axio Zoom.V16 (Zeiss, Göttingen, Germany). Flat-mounts were divided into 16 quadrants (200 µm x 200 µm) and Brn3a^+^ and Iba1^+^ cells were quantified. Groups were compared using one-way ANOVA followed by Tukey’s post hoc test. The WT CO group was set to 100 %.

### 2.7 Western blotting

Retinal tissue (n = 5/group) was homogenized in 150 µl and optic nerve tissue (n = 5/group) in 100 µl lysis buffer (60 mM n-octyl-β-D-glucopyranoside, 50 mM sodium acetate, 50 mM Tris chloride, pH 8.0 and 2 M urea) containing a protease inhibitor cocktail (Sigma-Aldrich) for 1 h on ice. Prior lysis, the optic nerve tissue was incubated in liquid nitrogen. Subsequently, all samples were centrifuged at 14.000 x g at 4°C for 30 minutes and the supernatant was used to determine the protein concentration. A BCA Protein Assay kit (Pierce, Thermo Fisher Scientific, Rockford, IL, USA) was used for retinal tissue. For optic nerves, the Qubit® Protein Assay kit (Life Technologies GmbH, Darmstadt, Germany) was used according to manufacturer’s instructions. 4x SDS buffer was added to each protein sample (20 µg) and denaturized for 5 minutes at 94°C. After separation via SDS-PAGE (10 % gels respectively 4–12 % polyacrylamide gradient gels), proteins were transferred to a polyvinylidene difluoride (PVDF) membrane (Roth, Karlsruhe, Germany) by Western blotting (1-2 h and 74 mA). Membranes were blocked (5 % w/v milk powder in TRIS-buffered saline (TBS) and Tween 20, TBST) at RT for 1 h and incubated with the primary antibody (Table 3) in blocking solution at 4°C overnight. Next day, membranes were washed with TBST and incubated with horseradish peroxidase (HRP) coupled secondary antibodies (Table 3) in blocking solution at RT for 2 h. Excess antibody was washed off with TBST. ECL Substrate (Bio-Rad Laboratories GmbH, München, Germany) was used to develop the membrane (mixed 1:1 for 5 minutes). Finally, protein immunoreactivity was detected with a MicroChemi Chemiluminescence Reader (Biostep, Burkhardtsdorf, Germany). Band intensity was analyzed using ImageJ software and normalized to a corresponding reference protein (β-Actin/vinculin). The normalized values of the Western blot results were given in arbitrary units (a.u.).

**Table 3:** List of primary and secondary antibodies for western blotting. Primary and secondary antibodies, including species, clonality, type, dilution, source and stock number/RRID are listed. Relative quantification of band intensity was done against the housekeeping proteins β-Actin or vinculin. HRP = horseradish peroxidase, kDa = Kilodalton.

| Primary Antibody | Species, Clonality, Type | Dilution | Source (Stock no.)/RRID | Secondary Antibody, Species, Type | Dilution/Source | kDa |
| --- | --- | --- | --- | --- | --- | --- |
| β-Actin | Mouse, monoclonal, IgG | 1:5000 | BD Bioscience (612657)<br>RRID:AB_399901 | Anti-mouse, IgG + IgM HRP | 1:5000/<br>Dianova | 42 |
| GFAP | Rabbit, polyclonal, IgG | 1:10000 | Dako (Z0334)<br>RRID:AB_10013382 | Anti-rabbit, IgG HRP | 1:5000/<br>Dianova | 50 |
| MBP | Rat, monoclonal, IgG | 1:1000 | Bio-Rad (MCA409)<br>RRID:AB_325004 | Anti-rat, IgG HRP | 1:5000/<br>Dianova | 20 |
| Olig2 | Rabbit, polyclonal, IgG | 1:500 | Merck (AB9610)<br>RRID:AB_570666 | Anti-rabbit, IgG HRP | 1:5000/<br>Dianova | 32 and 50 |
| vinculin | Mouse, monoclonal, IgG1 | 1:200 | Sigma-Aldrich (V 9131)<br>RRID:AB_477629 | Anti-mouse, IgG + IgM HRP | 1:5000/<br>Dianova | 116 |

### 2.8 RNA isolation, cDNA synthesis, and RT-qPCR

Retinae and optic nerves were explanted 10 weeks after immunization and stored at -80°C until purification (n = 5/group). The RNA isolation of the retina was carried out according to the manufacturer’s introduction using the Gene Elute Mammalian Total RNA Miniprep Kit (Sigma-Aldrich, St. Louis, MO, USA). For total RNA isolation of optic nerve tissue, the ReliaPrepTM RNA Tissue Miniprep System (Promega, Madison, WI, USA) was used. Prior isolation optic nerve tissue was incubated in liquid nitrogen. The concentration and purity of the isolated RNA was determined photometrically using the BioSpectrometer® (Eppendorf, Hamburg, Germany). 1 µg RNA and random hexamer primers were used for reverse transcription using the cDNA synthesis kit (Thermo Fisher Scientific, Waltham, MA, USA). RT-qPCR experiments were done with SYBR Green I in a Light Cycler 96® (Roche Applied Science, Mannheim, Germany). For each primer pair (Table 4) efficiencies were determined by a dilution series of 5, 25 and 125 ng cDNA. Expression in retina and optic nerve tissue was normalized against the housekeeping genes *β-Actin* (*Actb*) and *18S*, respectively.

**Table 4:**
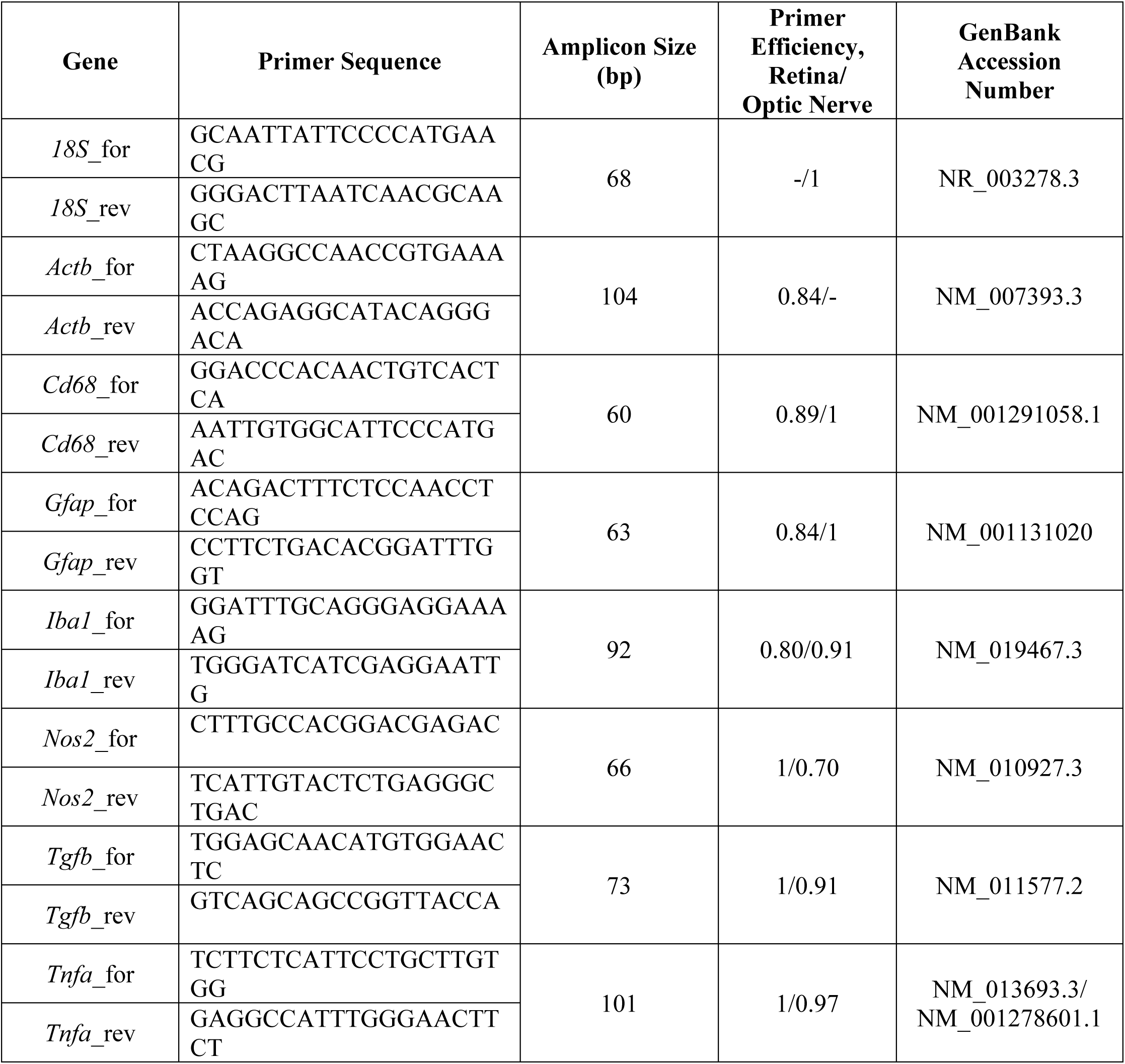
List of primer pairs used for mRNA analyses by RTq-PCR. For evaluation of mRNA-levels, *Actb* and *18S* served as housekeeping genes. Primer sequence, predicted amplicon size, primer efficiency for retinae and optic nerves, and GenBank accession number are listed. bp = base pairs, for = forward, rev = reverse.

### 2.9 Statistical analysis

Immunohistological, Western blot, IOP, and ERG data of control WT (WT CO) and KO (KO CO) as well as ONA-immunized WT (WT ONA) and KO (KO ONA) were analyzed by one-way ANOVA followed by Tukey’s post hoc test using Statistica software (V13.3; StatSoft (Europe), Hamburg, Germany). Results of IOP measurements were presented as mean ± standard error mean (SEM) ± standard deviation (SD). ERG recordings, immunohistochemical, and Western blot data were shown as mean ± SEM. For RT-qPCR results, groups were compared using the pairwise fixed reallocation and randomization test (REST© software) and were presented as median ± quartile ± minimum/maximum (Pfaffl et al., 2002).

## 3 Results

### 3.1 No changes in IOP and retinal functionality in the EAG mouse model

IOP measurements were performed before immunization in WT (WT CO) and KO (KO CO) at 5 weeks of age (Figure 1A). After immunization, IOP was measured weekly in control and immunized WT (WT ONA) and KO (KO ONA) animals until the end of the study. At 5 weeks of age (−1), we observed no significant differences in the IOP of WT CO (9.8 ± 0.2 mmHg) and KO CO (9.7 ± 0.1 mmHg; p = 1.0). Furthermore, no changes in the IOP were found in control and immunized groups throughout the study (Supplementary Table 1).

**Figure 1:**
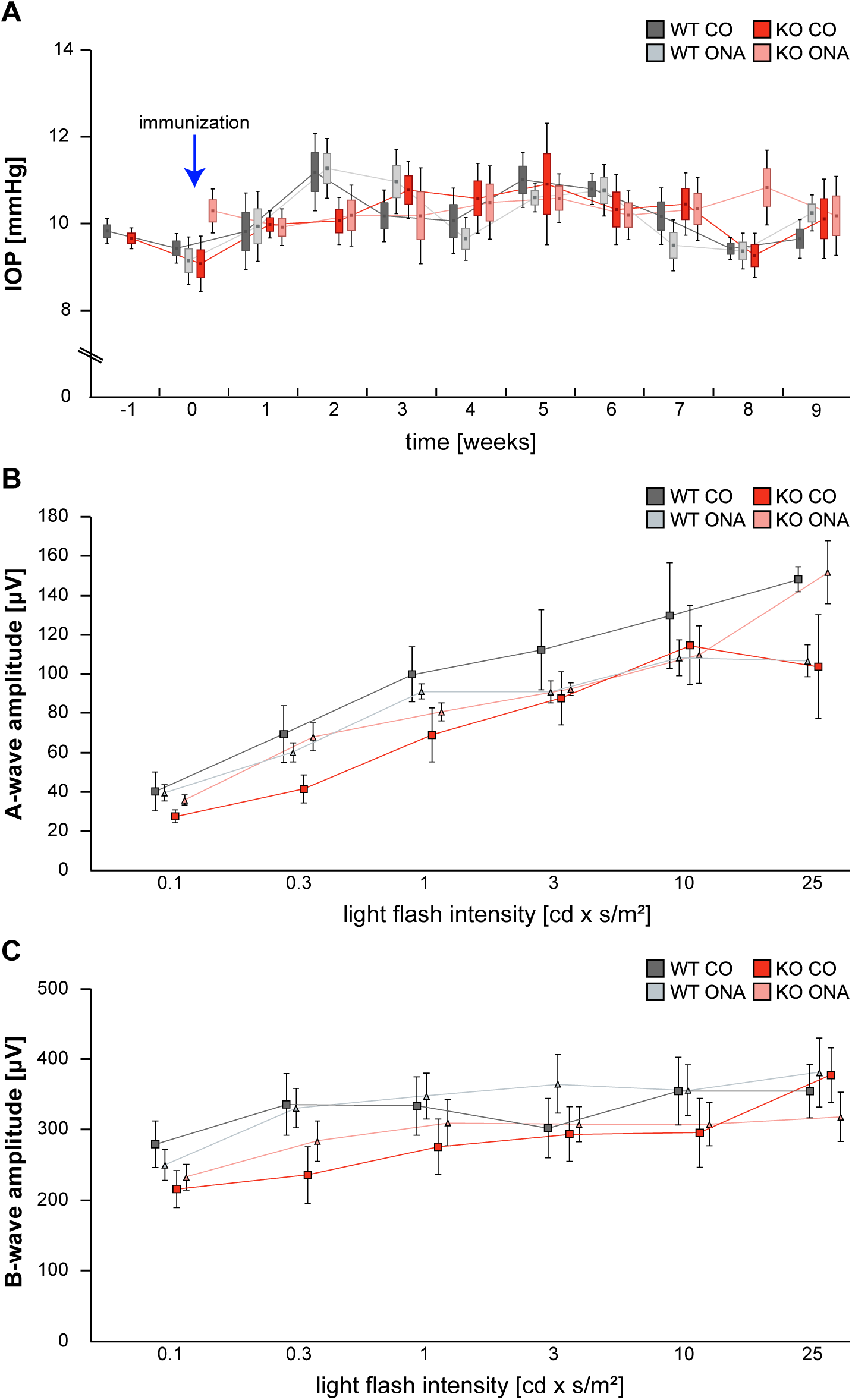
IOP and ERG recordings were not altered after immunization of WT and KO mice. **(A)** IOP measurements were performed before immunization in WT and KO at 5 weeks of age (−1; n = 16/group). Then, IOP was determined weekly in immunized and control WT and KO until the end of the study (n = 8/group). No significant changes could be detected between all groups (n = 5/group). **(B, C)** ERG recordings 10 weeks after initial immunization in control and immunized WT and KO mice. No changes in a-wave **(B)** and b-wave **(C)** amplitudes could be detected in control and immunized WT and KO mice. Data were analyzed using one-way ANOVA followed by Tukey’s post hoc test and present as means ± SEM ± SD in **(A)** and mean ± SEM in **(B, C)**. cd: candela; IOP: intraocular pressure; µV: micro volt; m: minutes; s: seconds.

To determine possible retinal function deficits, induced by ONA-immunization, we performed ERG recordings of control and immunized WT and KO mice. Under scotopic conditions, a-wave responses arise from rod-photoreceptors, while b-waves represent the rod bipolar and Müllerglia cell response. In all four conditions no significant differences were observed between control and immunized WT and KO animals (Figure 1B, C; Supplementary Table 2). Therefore, we concluded that photoreceptor and bipolar cell function was not affected in this EAG mouse model.

### 3.2 Significant loss of RGCs following immunization

Previous studies of an EAG rat model showed a significant reduction of RGCs 4 weeks post immunization with ONA (Laspas et al., 2011; Noristani et al., 2016). Additionally, an upregulation of Tnc was found before significant loss of RGCs (Reinehr et al., 2016). Based on these findings, immunohistochemical stainings of RGCs were performed with an antibody against Brn3a, which specifically detects RGCs (Figure 2, Supplementary Table 3). The evaluation of RGCs in retinal cross-sections showed a significant reduction in the percentage of Brn3a^+^ cells in WT ONA compared to WT CO as well as to KO CO (WT ONA: 73.1 ± 6.1 % Brn3a^+^ cells vs. WT CO: 100.0 ± 4.2 % Brn3a^+^ cells; p = 0.004 and KO CO: 92.2 ± 3.9 % Brn3a^+^ cells, p = 0.04, Figure 2A, B). Interestingly, no significant differences between control and immunized KO could be detected in horizontal cross-sections (KO CO: 92.2 ± 3.9 % Brn3a^+^ cells vs. KO ONA: 83.7 ± 8.7 % Brn3a^+^ cells, p = 0.57, Figure 2A, B).

**Figure 2:**
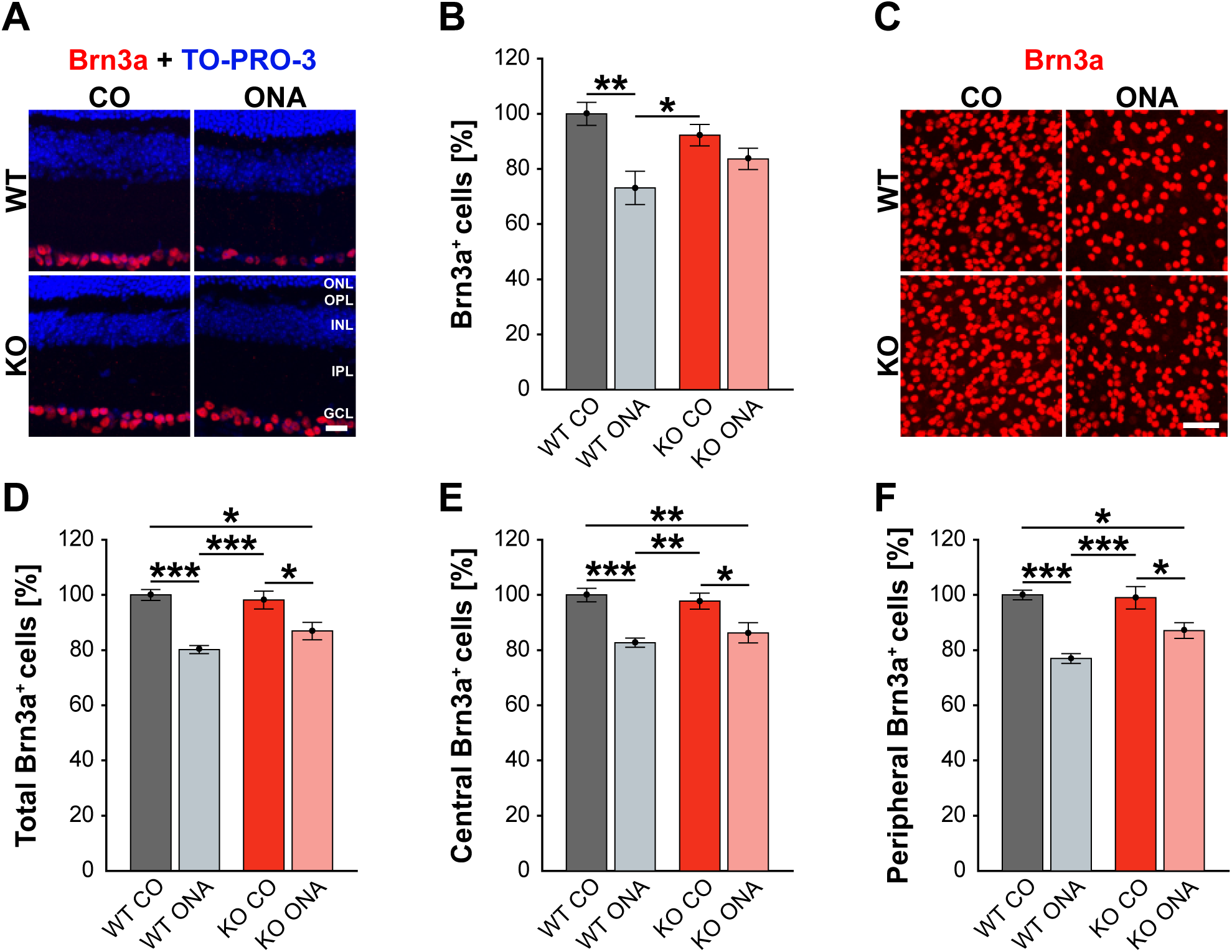
Lower RGC loss in immunized KO animals. **(A)** Retinal cross-sections from WT CO, WT ONA, KO CO, and KO ONA mice were stained with an antibody against Brn3a (red), and nuclei were detected with TO-PRO-3 (blue). **(B)** A decline of RGC density was detected in WT ONA compared to the control groups (n = 5/group). **(C)** Representative pictures of Brn3a^+^ cells in retinal flat-mounts. **(D-F)** Quantification of the total RGC number as well as in central and peripheral parts (n = 9/group). A significant loss of RGCs was detected in immunized WT and KO in comparison to the control groups. It was also shown that the RGC number in KO ONA RGCs were significantly decreased compared to WT CO. Furthermore, WT ONA RGC number were significantly reduced compared to KO CO. Data were analyzed using one-way ANOVA followed by Tukey’s post hoc test and values were shown as mean ± SEM. *p < 0.05; **p < 0.01; ***p < 0.001. Scale bar = 20 µm in **(A)** and 50 µm in **(C)**. ONL: outer nuclear layer; OPL: outer plexiform layer; INL: inner nuclear layer; IPL: inner plexiform layer; GCL: ganglion cell layer.

To further characterize the RGC population, we counted Brn3a^+^ cells in retinal flat-mounts (Figure 2C). We determined the total amount (Figure 2D) as well as the number of Brn3a^+^ cells within the central (Figure 2E) and peripheral (Figure 2F) area of the retina. A significant reduction in the total number was observed in immunized WT compared to both control genotypes (WT ONA: 80.3 ± 1.5 % Brn3a^+^ cells vs. WT CO: 100.0 ± 2.0 % Brn3a^+^ cells, p < 0.001, KO CO: 98.2 ± 3.3 % Brn3a^+^ cells, p < 0.001). Also, a significant loss of RGCs was detected in KO ONA mice (86.9 ± 3.1 % Brn3a^+^ cells) compared to KO CO (p = 0.02) and WT CO (p = 0.01). A comparable percentage of Brn3a^+^ cells was also observed in the central retina. So, immunized WT and KO animals showed a significant decline of RGCs compared to the corresponding control groups (WT ONA: 82.7 ± 1.7 % Brn3a^+^ cells vs. WT CO: 100.0 ± 2.5 % Brn3a^+^ cells, p < 0.001 and KO ONA: 86.3 ± 3.7 % Brn3a^+^ cells vs. KO CO: 97.8 ± 2.9 % Brn3a^+^ cells, p = 0.03). No significant differences were found between both immunized genotypes (p = 0.80). Furthermore, a decrease in the RGC density was verified in the peripheral area. Retinae of the WT ONA group (77.0 ± 1.8 % Brn3a^+^ cells) displayed a loss of about 25 % RGCs compared to WT CO (100.0 ± 1.7 % Brn3a^+^ cells, p < 0.001). A significant reduction was also found in the comparison of KO CO and KO ONA (KO CO: 99.0 ± 4.1 % Brn3a^+^ cells vs. KO ONA: 87.1 ± 2.8 % Brn3a^+^ cells, p = 0.02). However, KO ONA showed a decrease of about 15 % in the peripheral part compared to WT CO group (p = 0.01).

Collectively, we found a weaker RGC damage in immunized KO and the loss of RGCs was more prominent in the WT condition.

### 3.3 Optic nerve degeneration post ONA-immunization in WT mice

To analyze a possible degeneration of RGC axons, immunoreactivity of βIII-tubulin was examined in optic nerve longitudinal sections of control and immunized WT and KO animals (Figure 3A). The immunopositive area of βIII-tubulin was significantly reduced in immunized WT (30.85 ± 8.55 % βIII-tubulin^+^ area) compared to WT CO (100.00 ± 18.35 % βIII-tubulin^+^ area, p = 0.04, Figure 3B). Also, the βIII-tubulin^+^ area was decreased compared to both KO conditions (KO CO: 93.66 ± 19.18 % βIII-tubulin^+^ area, p = 0.06 and KO ONA: 119.93 ± 13.48 % βIII-tubulin^+^ area, p < 0.01).

**Figure 3:**
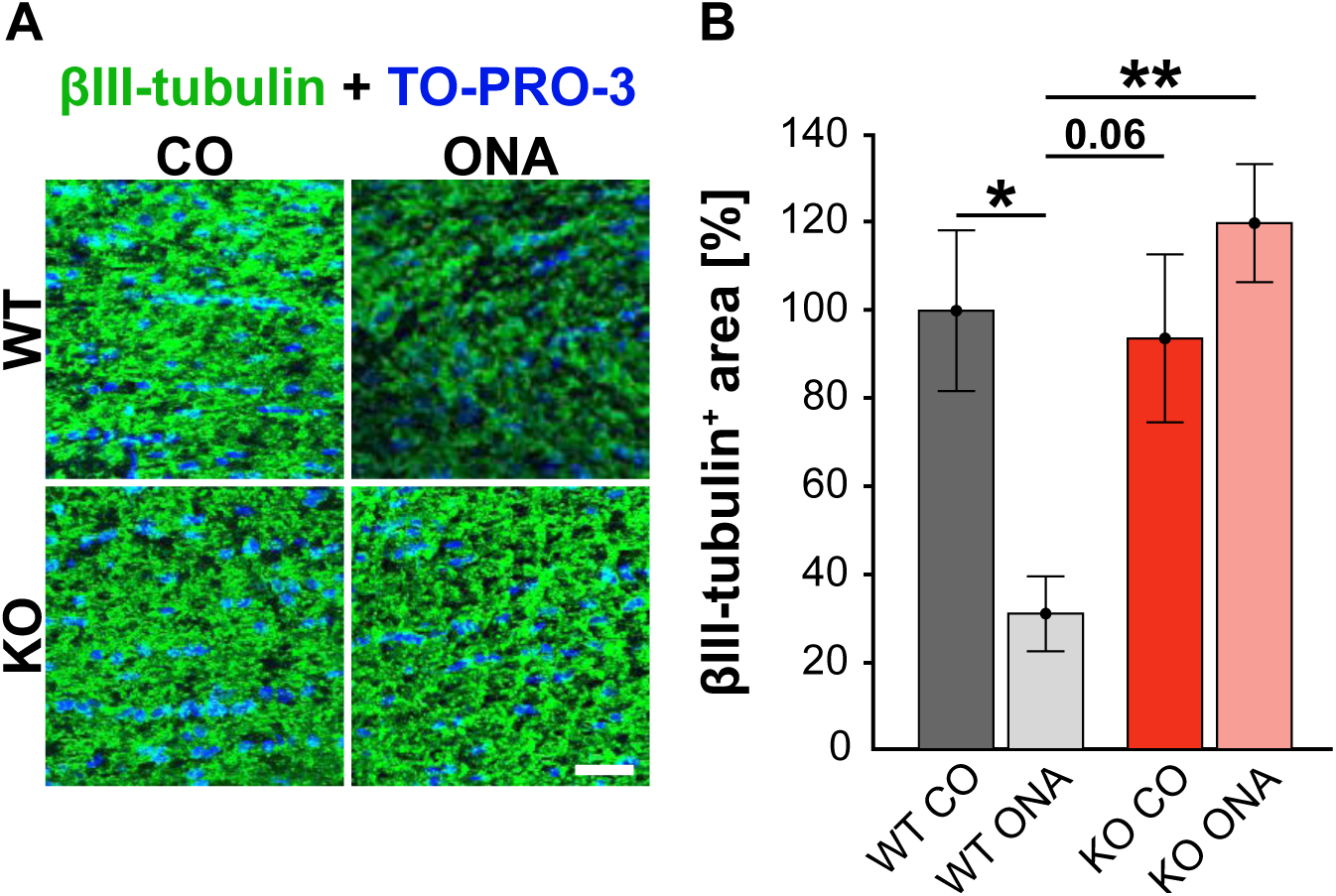
No optic nerve degeneration in KO post immunization. **(A)** Optic nerve slices were stained with βIII-tubulin (green) and cell nuclei were marked with TO-PRO-3 (blue). **(B)** WT ONA mice showed a significantly decreased βIII-tubulin^+^ area compared to WT CO as well as to KO ONA group. Data were analyzed using one-way ANOVA followed by Tukey’s post hoc test and present as means ± SEM. *p < 0.05; **p < 0.01. n = 4/group. Scale bar = 20 µm.

### 3.4 Extenuated macroglial reactivity after immunization in KO mice

Our results showed a decrease in the RGC number 10 weeks after immunization in WT and KO animals. Next, we investigated, if this glaucomatous neurodegeneration is associated with an altered macroglial response. Therefore, we analyzed the immunoreactivity of GFAP^+^ astrocytes in retinal cross-sections. GFAP stained astrocytes were localized in the ganglion cell layer (GCL; Figure 4A). The GFAP^+^ area was increased in WT ONA (167.22 ± 18.61 % GFAP^+^ area) compared to the control groups (WT CO: 100.00 ± 9.28 % GFAP^+^ area, p < 0.05 and KO CO: 105.81 ± 4.54 % GFAP^+^ area, p = 0.02, Figure 4B). Interestingly, no changes in the GFAP signal were found in KO ONA (142.49 ± 8.19 % GFAP^+^ area) compared to KO CO (p = 0.18) as well as to WT CO (p = 0.11). The statistical comparison of both immunized genotypes showed no significant differences (p = 0.46).

**Figure 4:**
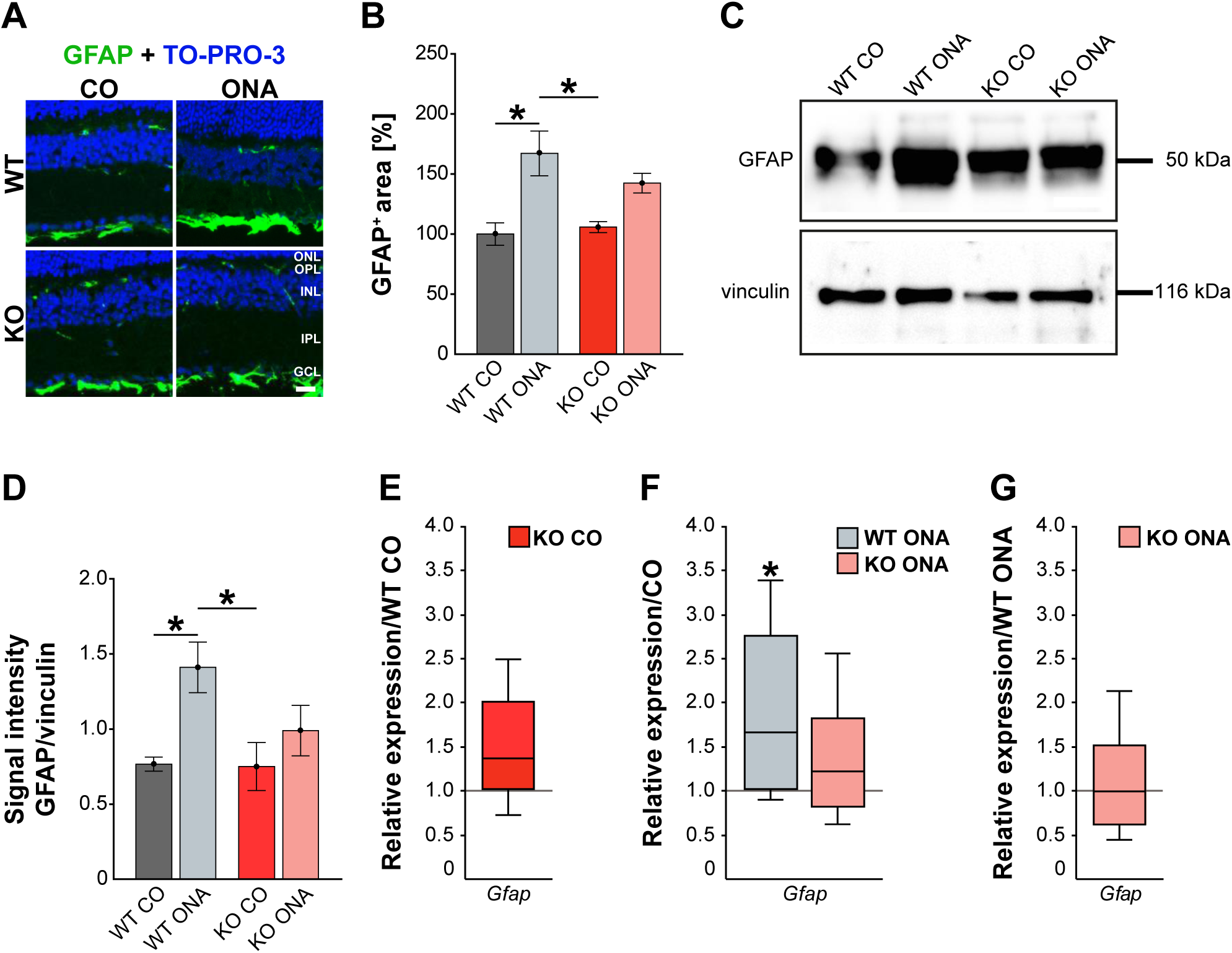
Reduced astrogliosis after immunization in KO mice. **(A)** Images of GFAP stained macroglia cells in retinal cross-section from control and immunized WT and KO animals. Immunohistochemistry revealed a prominent signal for GFAP^+^ cells (green) in the GCL and NFL. Cell nuclei were detected with TO-PRO-3 (blue). **(B)** GFAP^+^ area was significantly increased in WT ONA compared to WT CO. No changes of the GFAP signal area was found in KO ONA compared to the control groups. **(C)** Western blot analyses of GFAP protein in retinal tissue. **(D)** Quantification revealed more GFAP in WT ONA, whereas a comparable level was observed in KO ONA. **(E)** No differences of *Gfap* expression were noted in KO CO compared to WT CO. **(F)** A significant upregulation of *Gfap* mRNA expression was seen in WT ONA in comparison to WT CO. **(G)** WT ONA and KO ONA animals showed similar *Gfap* levels. Data were analyzed using one-way ANOVA followed by Tukey’s post hoc test and present as means ± SEM in **(B, D)**. For RT-qPCR results, groups were compared using the pairwise fixed reallocation and randomization test and were shown as median ± quartile ± minimum/maximum in **(E-G)**. *p < 0.05. n = 5/group. Scale bar = 20 µm. ONL: outer nuclear layer, OPL: outer plexiform layer, INL: inner nuclear layer, IPL: inner plexiform layer, GCL: ganglion cell layer.

Then, we evaluated GFAP protein levels via Western blot. For GFAP, a prominent band was detected at 50 kDa (Figure 4C). Relative quantification verified a significant increase in the GFAP protein concentration in WT after immunization (WT ONA: 1.41 ± 0.17 a.u. vs. WT CO: 0.77 ± 0.11 a.u., p = 0.03 and KO CO: 0.75 ± 0.16 a.u., p = 0.03, Figure 4D). No changes were observed in the GFAP level of control and immunized KO animals (KO ONA: 0.98 ± 0.16 a.u., p

= 0.66).

We also analyzed the mRNA expression of *Gfap* in retinae via RT-qPCR (Figure 4E-G, Supplementary Table 4). Analysis revealed comparable levels of *Gfap* in KO CO and WT CO (1.4-fold, p = 0.11, Figure 4E). A significant increase of *Gfap* mRNA expression levels was observed in WT ONA (WT CO vs. WT ONA: 1.7-fold, p = 0.04, Figure 4F), whereas no differences could be detected in KO ONA (KO CO vs. KO ONA: 1.2-fold, p = 0.4, Figure 4F). The expression was comparable in both immunized genotypes (1.0-fold, p = 0.99, Figure 4G).

For GFAP, a thread-like staining pattern could be observed in optic nerve slices (Figure 5A). The evaluation of the GFAP immunoreactivity in optic nerve sections also showed no increased macroglial area in KO post immunization (KO ONA: 134.30 ± 23.57 % GFAP area vs. KO CO: 147.18 ± 31.27 % GFAP^+^ area, p = 0.98 and WT CO: 100.00 ± 17.72 % GFAP^+^ area, p = 0.70, Figure 5B). Moreover, a nearly doubled GFAP intensity was observed in WT ONA (196.70 ± 13.60 % GFAP^+^ area) compared to the corresponding control group (p = 0.04).

**Figure 5:**
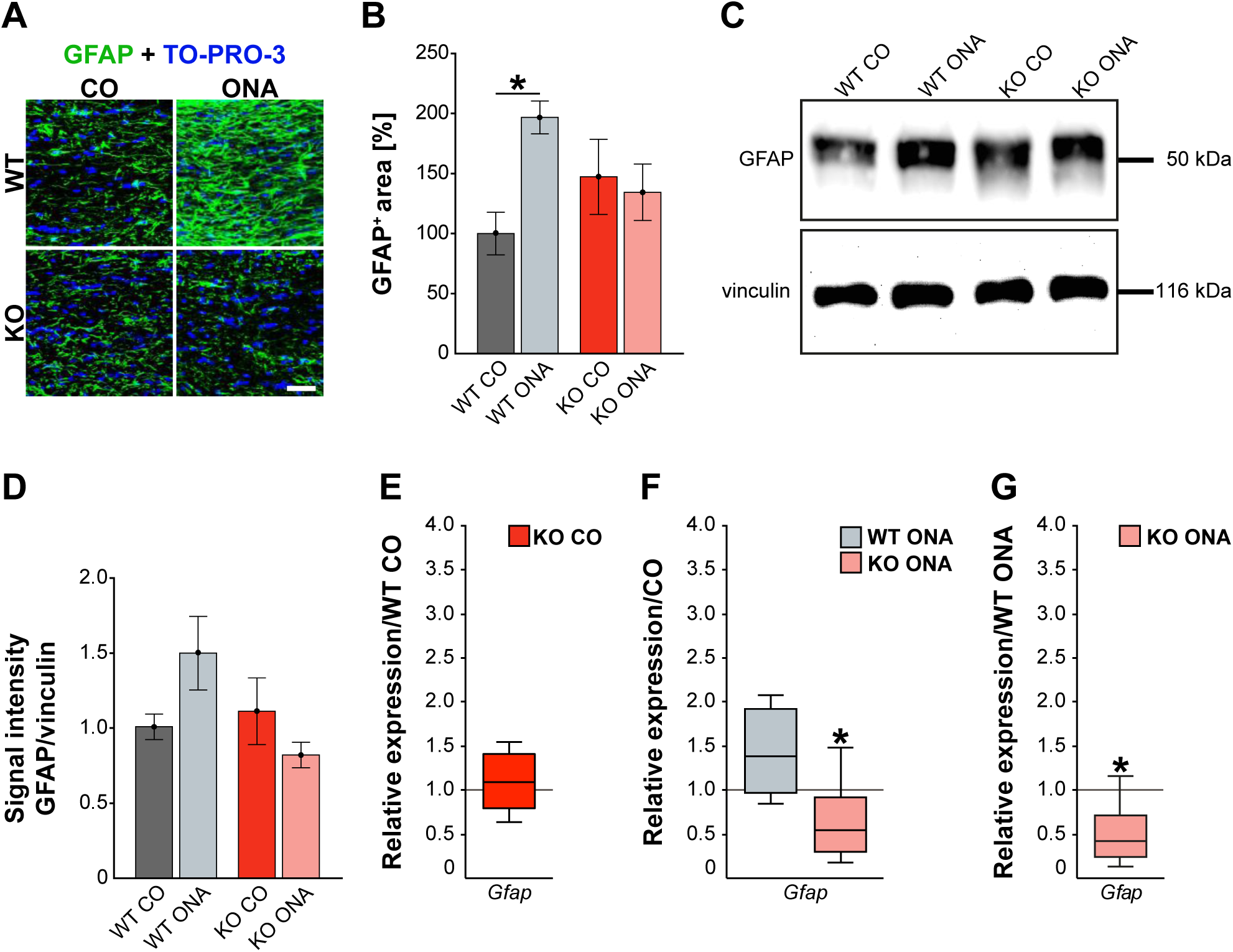
Diminished macroglial response post ONA-immunization in optic nerve tissue of KO animals. **(A)** Representative pictures of optic nerve sections of control and immunized WT and KO mice stained against GFAP (green). Nuclear staining was done with TO-PRO-3 (blue). **(B)** WT ONA animals showed a larger GFAP^+^ staining area then WT CO. No differences between KO mice could be detected. **(C)** Western blot analyses of relative GFAP protein levels in optic nerve tissue. **(D)** Protein quantification revealed slightly enhanced band intensity in WT ONA, whereas KO ONA exhibit no increased GFAP level. **(E)** No differences of *Gfap* expression were noted in KO CO compared to WT CO. **(F)** A slight upregulation of *Gfap* mRNA expression was seen in WT ONA in comparison to WT CO, but in KO ONA decreased expression was verified compared to KO CO. **(G)** A downregulation of *Gfap* in KO ONA in comparison to WT ONA was noted. Data were analyzed using one-way ANOVA followed by Tukey’s post hoc test and were shown as mean ± SEM in **(B, D)**. For RT-qPCR, groups were compared using the pairwise fixed reallocation and randomization test and were shown as median ± quartile ± minimum/maximum in **(E-G)**. *p < 0.05. n = 5/group. Scale bar = 20 µm.

Furthermore, protein levels of GFAP via Western blot analyses were comparable between all four groups (Figure 5C). However, the band intensity in WT ONA group was tendentially increased compared to the control group (WT ONA: 1.50 ± 0.25 a.u. vs. WT CO: 1.01 ± 0.08 a.u., p = 0.24, Figure 5D). Equal protein levels were found between control and ONA KO animals (KO CO: 1.11 ± 0.22 a.u. vs. KO ONA: 0.82 ± 0.08 a.u., p = 0.65).

Finally, the RT-qPCR results of the optic nerve tissue showed no changes in *Gfap* expression between both control groups (WT CO vs. KO CO: 1.1-fold, p = 0.54, Figure 5E, Supplementary Table 4). In line with the immunohistochemical results, we found a slightly enhanced mRNA level in WT ONA (WT CO vs. WT ONA: 1.4-fold, p = 0.07, Figure 5F), whereas the KO ONA animals exhibited a reduction of *Gfap* expression (KO CO vs. KO ONA: 0.5-fold, p < 0.05). Interestingly, in a direct comparison of the two immunized groups, *Gfap* expression was significantly reduced in KO animals (WT ONA vs. KO ONA: 0.4-fold, p = 0.02, Figure 5G). In summary, we concluded that Tnc deficiency resulted in a diminished macroglial reaction during retinal and optic nerve degeneration in the EAG mouse model.

### 3.5 Decreased demyelination after ONA-immunization in KO mice

Our study demonstrates RGC degeneration in WT and KO animals after immunization. Furthermore, we noted that Tnc deficiency resulted in a diminished macroglial response. Finally, we analyzed the impact of ONA-immunization on oligodendroglia in optic nerve tissue. The oligodendrocytes appear in two different populations, as immature oligodendrocytes precursor cells (OPCs) and as myelinating, mature oligodendrocytes. To analyze both oligodendrocyte populations separately, an immunohistochemical colocalization staining was performed using the markers Olig2 and CC1. Olig2 is expressed by oligodendrocytes of all stages (Gautier et al., 2015). In contrast, CC1 is only expressed by mature oligodendrocytes (Bin et al., 2016). Colocalization identified double positive cells as mature and single Olig2^+^ cells as immature oligodendrocytes. In addition, we also investigated MBP on protein level via immunohistochemistry and Western blot. This protein is specifically expressed by myelinating oligodendrocytes (Pohl et al., 2011). Immunohistochemical stainings revealed fewer Olig2^+^ cells in the WT ONA group compared to the other groups. Interestingly, there were more Olig2^+^ cells in KO ONA than in WT ONA (Figure 6A). 76.3 ± 1.6 % Olig2^+^ cells were found in WT ONA, which indicates a significant oligodendrocyte loss over 25 % compared to control WT (100.0 ± 3.5 % Olig2^+^ cells, p < 0.001, Figure 6B). The number of Olig2^+^ cells was also significantly decreased in WT ONA compared to KO CO (p = 0.04). No differences were observed between both Tnc deficient groups (KO CO: 90.2 ± 2.4 % Olig2^+^ cells vs. KO ONA: 95.2 ± 4.9 % Olig2^+^ cells, p = 0.71). Most interestingly, we verified significant differences between both immunized groups (p < 0.01). The number of double-positive (Olig2^+^/CC1^+^) mature oligodendrocytes was clearly reduced in WT ONA compared to all other groups (Figure 6A). The statistical evaluation demonstrated only 64.7 ± 5.8 % Olig2^+^/CC1^+^ cells in WT ONA, whereas KO ONA exhibited 107.8 ± 8.9 % Olig2^+^/CC1^+^ cells in optic nerve slices (p = 0.002, Figure 6C). Also, immunized WT showed a significant loss of mature oligodendrocytes compared to WT CO (100.0 ± 8.3 % Olig2^+^/CC1^+^ cells, p = 0.01) and KO CO (118.4 ± 3.5 % Olig2^+^/CC1^+^ cells, p < 0.001).

**Figure 6:**
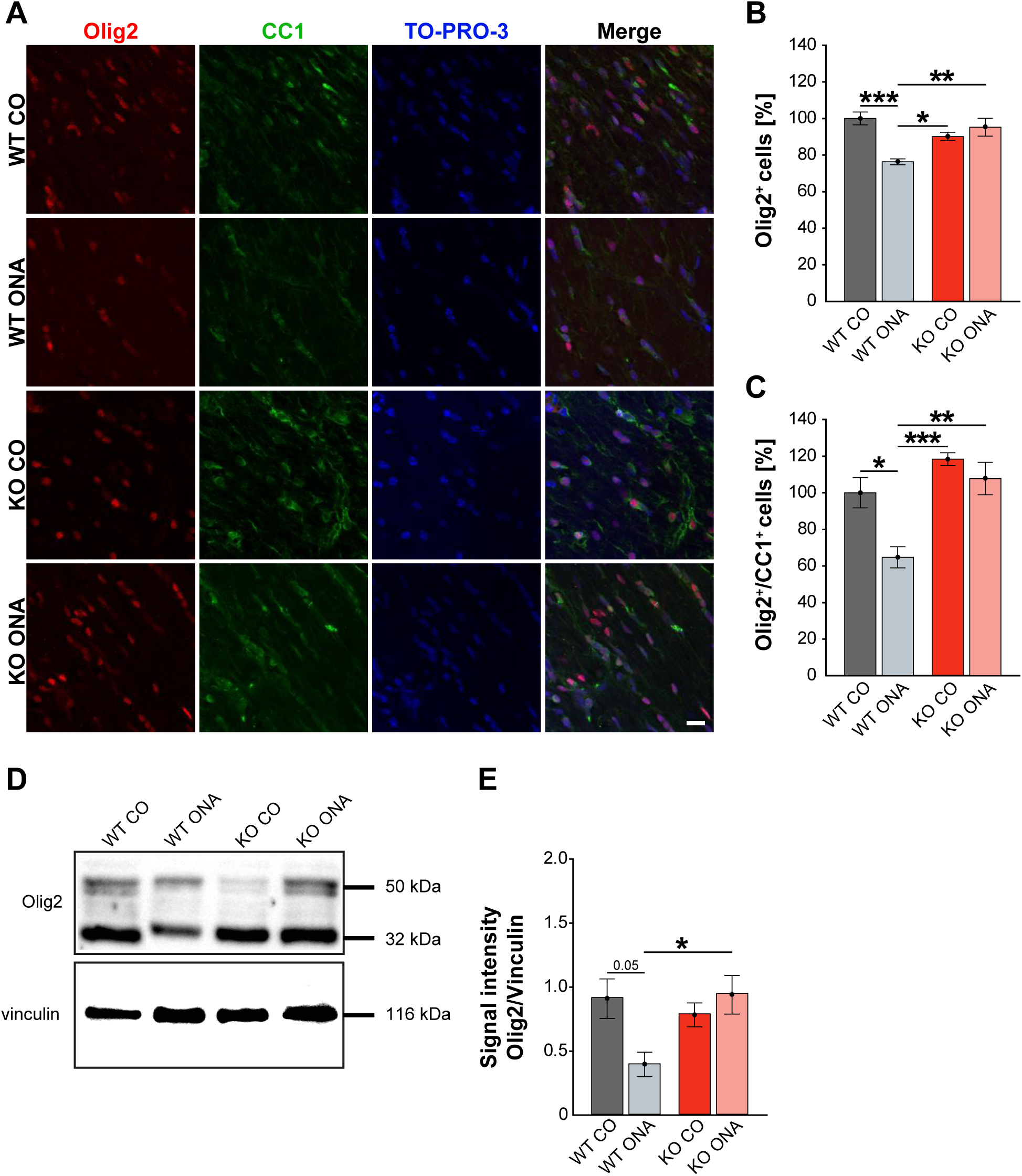
No demyelination after immunization in KO mice. **(A)** Olig2 (red) and CC1 (green) staining of optic nerve sections. Cell nuclei were labeled with TO-PRO-3 (blue). **(B)** Quantification of Olig2^+^ cells revealed a significant decrease of oligodendroglia in WT ONA compared to WT CO and KO CO. Interestingly, the statistical comparison of both immunized groups showed a significant loss of Olig2^+^ cells in WT compared to KO mice. **(C)** WT ONA nerves displayed a significantly decrease of mature oligodendrocytes in comparison to both control groups. A significant higher amount of double positive oligodendrocytes was observed in KO ONA compared to WT ONA. **(D)** An exemplary Western blot of Olig2. **(E)** Relative protein quantification revealed a slightly enhanced band intensity of the Olig2 protein in WT ONA, whereas KO ONA nerves exhibited no reduction of the Olig2 protein level. Data were analyzed using one-way ANOVA followed by Tukey’s post hoc test and values were shown as mean ± SEM. *p < 0.05; **p < 0.01, ***p < 0.001. n = 5/group. Scale bar = 50 µm.

To consolidate the immunohistochemistry results, we analyzed the Olig2 protein level in optic nerves by Western blot analyses (Figure 6D). For Olig2, we observed two bands at 32 kDa and 50 kDa. A decrease of the band intensity was found in WT ONA (0.38 ± 0.21 a.u.) compared to the corresponding control group (0.89 ± 0.35 a.u., p = 0.05, Figure 6E). Equal Olig2 protein levels were observed in KO CO (0.77 ± 0.21 a.u.) and KO ONA (0.92 ± 0.34 a.u., p = 0.82). Interestingly, missing Tnc resulted in a significantly higher Olig2 protein level post immunization compared to WT (p = 0.04).

Finally, myelinating oligodendrocytes were detected with an antibody against MBP. Immunohistochemical staining revealed significantly reduced MBP immunoreactivity in WT ONA (Figure 7A). Statistical analyses showed a decreased MBP signal in WT ONA (46.22 ± 11.30 % MBP^+^ area) compared to WT CO (100.00 ± 7.47 % MBP^+^ area, p = 0.001), KO CO (88.76 ± 4.93 % MBP^+^ area, p = 0.01) and KO ONA (109.79 ± 7.01 % MBP^+^ area p < 0.001, Figure 7B).

**Figure 7:**
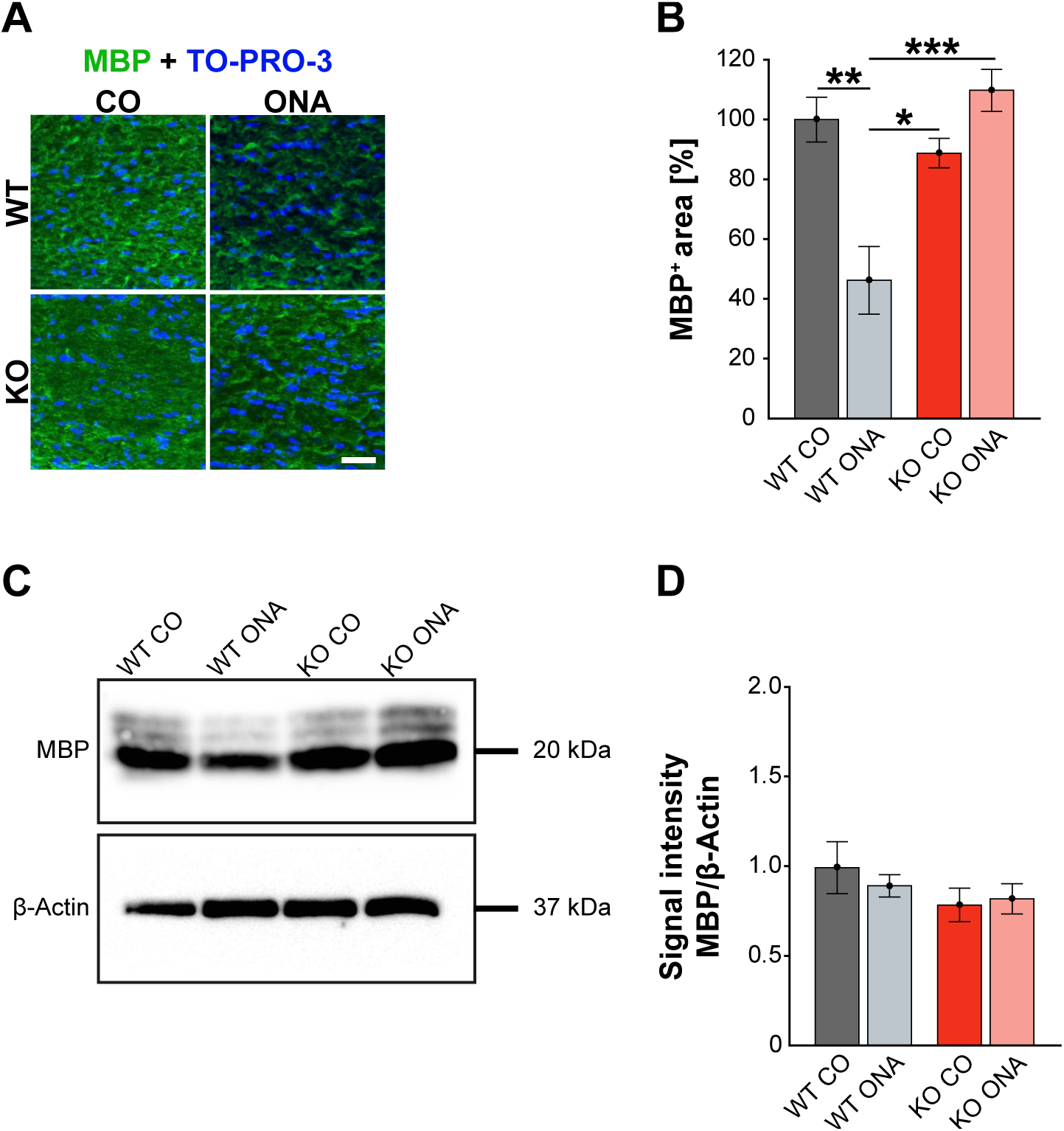
Unaltered MBP immunoreactivity post immunization in KO mice. **(A)** MBP (green) was stained in optic nerve tissue. In blue TO-PRO-3 detected cell nuclei. Immunohistochemistry indicates a reduced MBP signal in WT ONA. **(B)** A significant downregulation of MBP was noted in WT ONA compared to WT CO. Furthermore, the MBP signal was significantly reduced in WT ONA compared to control as well as to immunized KO mice. **(C)** Western blot analyses of MBP of optic nerve tissue. **(D)** Comparable MBP protein levels were observed in all groups. Data were analyzed using one-way ANOVA followed by Tukey’s post hoc test and values were indicated as mean ± SEM. *p < 0.05; **p < 0.01; ***p < 0.001. n = 5/group. Scale bar = 20 µm.

MBP was examined on protein level via Western blot analyses and a prominent protein band was detected at 20 kDa (Figure 7C). Quantitative analyses revealed comparable MBP protein levels in control (WT CO: 0.99 ± 0.14 a.u., KO CO: 0.82 ± 0.08 a.u.) and ONA mice (WT ONA: 0.89 ± 0.06 a.u., KO ONA: 0.82 ± 0.08 a.u., Figure 7D).

In conclusion, we found a significant decrease in mature as well as immature oligodendroglia in WT after immunization. Remarkably, immunized Tnc deficient mice showed no demyelination.

### 3.6 Decreased number of microglia and declined microglial response in KO ONA mice

Neurodegeneration is often accompanied by reactive microgliosis. In order to analyze the microglia population in the EAG mouse model and the effects of immunization, we performed immunohistochemical staining of retinal flat-mounts using an Iba1 antibody (Figure 8A, Supplementary Table 3). The number of Iba1^+^ cells in the total as well as in the central and peripheral area of the retina was evaluated (Figure 8B-D). A significant increase in microglia numbers was detected in WT ONA (123.0 ± 2.4 % Iba1^+^ cells) compared to control WT and KO in the total retina (WT CO: 100.0 ± 2.9 % Iba1^+^ cells, p < 0.001 and vs. KO CO: 102.3 ± 5.7 % Iba1^+^ cells; p = 0.002, Figure 8B). No differences were found between both control groups (p = 0.97). Remarkably, a significantly lower number of Iba1^+^ cells was observed after immunization in KO ONA (KO ONA: 84.5 ± 2.7 % Iba1^+^ cells) compared to WT ONA (p < 0.001), WT CO (p = 0.03), and KO CO (p < 0.01). Also, in the central part of the retina 20 % more Iba1^+^ cells were detected in WT ONA (122.5 ± 2.9 % Iba1^+^ cells) compared to the corresponding control group (WT CO: 100.0 ± 3.5 % Iba1^+^ cells, p < 0.01, Figure 8C). Immunized WT also showed significantly more microglial cells compared to control (p < 0.001) and immunized KO mice (p < 0.001). In KO CO, we counted 98.1 ± 5.8 % Iba1^+^ cells, whereas KO ONA only has 84.0 ± 2.9 % Iba1^+^ cells (p = 0.08). A reduced microglia number was noted in KO ONA compared to WT CO (p = 0.04). Equal numbers of microglial cells were seen in both control groups (p = 0.99). Similarly, the number of microglia in WT ONA (123.6 ± 3.4 % Iba1^+^ cells) was significantly enhanced compared to WT CO (100.0 ± 3.2 % Iba1^+^ cells, p = 0.002, Figure 8D) and KO CO (107.2 ± 6.4 % Iba1^+^ cells, p < 0.05) in peripheral regions of the retinal. Also, the number of Iba1^+^ cells was lower in KO ONA (85.1 ± 3.1 % Iba1^+^ cells) compared to KO CO (p < 0.01) and WT CO (p = 0.08) in the periphery. Additionally, a significantly reduced microglial response was detected in immunized KO and WT animals (p < 0.001). Regarding the quantification of Iba1^+^ cells in the periphery, both control groups had similar cell counts (p = 0.63).

**Figure 8:**
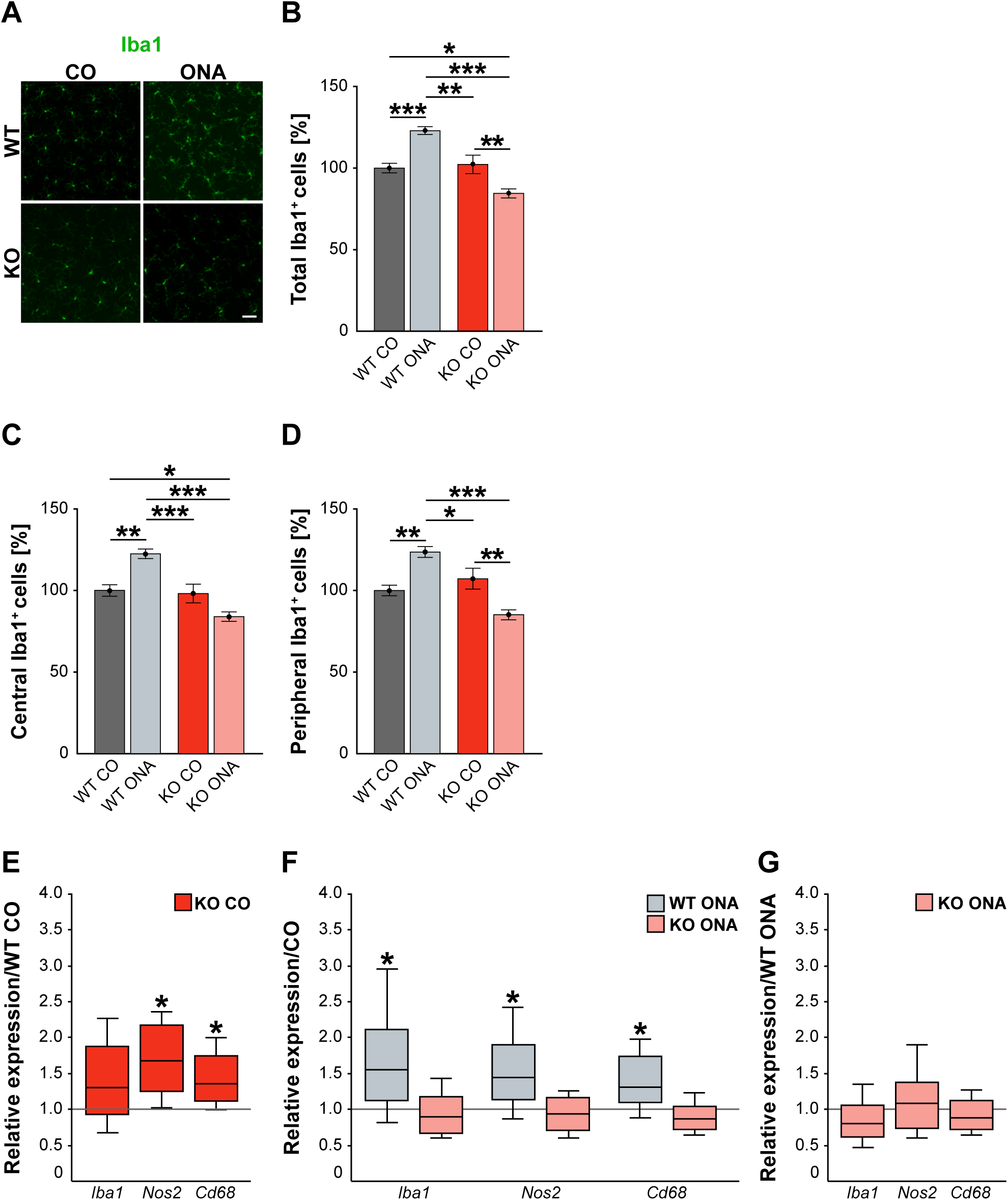
Decreased microglia response after immunization in KO mice. **(A)** Representative pictures of Iba1^+^ cells (green) in retinal flat-mounts of immunized and non-immunized WT and KO mice. **(B-D)** Quantification of Iba1^+^ microglia in control and immunized WT and KO animals in the total, central, and peripheral retina (n = 9/group). WT ONA group exhibited clearly more microglia. In contrast, KO ONA animals displayed fewer Iba1^+^ cells. **(E)** RT-qPCR analyses (n = 5/group) of the relative *Iba1, Nos2*, and *Cd68* mRNA expression showed a significant increase of *Nos2* and *Cd68* in KO CO compared to WT CO retinae. No differences were observed for *Iba1* mRNA expression. **(F)** Compared to WT CO, a significant upregulation of *Iba1, Nos2*, and *Cd68* levels were found in WT ONA. No significant changes were detected regarding the expression levels of these markers in KO ONA compared to KO CO. **(G)** After immunization, a comparable mRNA-Level of *Iba1, Cd68*, and *Nos2* was detected in KO ONA compared to KO CO. Data were analyzed using one-way ANOVA followed by Tukey’s post hoc test and present as means ± SEM in **(B-D)**. Groups were compared using the pairwise fixed reallocation and randomization test and were shown as median ± quartile ± minimum/maximum for mRNA analysis in **(E-G)**. *p < 0.05; **p < 0.01; ***p < 0.001. Scale bar = 50 µm.

In the next step, RT-qPCR was used to investigate whether microglia have reactive phenotypes. Beside *Iba1*, we also examined the markers *Nos2* and *Cd68* in retinal tissue (Figure 8E-G, Supplementary Table 4). No differences could be detected in the *Iba1* expression between control WT and KO mice (1.3-fold, p = 0.2, Figure 8E). However, KO CO mice showed significantly elevated levels of *Nos2* (1.7-fold; p = 0.013) and *Cd68* (1.3-fold; p = 0.017) compared to WT CO. The comparison of immunized and non-immunized WT showed a significantly increased expression of *Iba1* (1.5-fold, p = 0.048) as well as of the reactive markers *Nos2* (1.4-fold, p = 0.021) and *Cd68* (1.3-fold, p = 0.032, Figure 8F). Interestingly, comparable expression levels of these microglial markers were found in immunized KO and KO CO (p > 0.05, Figure 8F). RT-qPCR analyses revealed comparable mRNA levels in KO ONA compared to WT ONA (p > 0.05, Figure 8G).

In line with the RT-qPCR results of retinal tissue, we found a similar expression pattern of microglial markers in the optic nerve of control and immunized WT and KO mice (Supplementary Table 4 and Figure S1).

In summary, WT ONA animals showed a significantly increased microglia infiltration and glial marker expression, indicating an increased microglial response. Remarkably, a significantly reduced invasion and reactivity of microglia were observed in KO ONA, suggesting that Tnc signaling is an important modulator of microglia in glaucomatous neurodegeneration.

### 3.7 Altered expression pattern of pro- and anti-inflammatory cytokines in immunized WT compared to immunized KO animals

In our study, we noted a reactive gliosis and an increased microglial response after immunization in WT mice. Interestingly, these effects could not be detected in Tnc deficient animals. Next, we analyzed the pro- and anti-inflammatory responses of the microglial phenotypes in retinae and optic nerves (Figure 9, Supplementary Table 4). Here, *Tnfa* was used to study M1 pro-inflammatory microglia, while *Tgfb* is expressed by M2 anti-inflammatory microglia. RT-qPCR experiments revealed comparable mRNA level of *Tnfa* (1.4-fold, p = 0.132) and *Tgfb* (1.4-fold, p = 0.071) in KO CO mice compared to WT CO mice (Figure 9A). After ONA-immunization, *Tnfa* was significantly upregulated in WT and interestingly downregulated in KO compared to the corresponding control groups (WT CO vs. WT ONA: 1.7-fold, p = 0.026 and KO CO vs. KO ONA: 0.4-fold, p = 0.031, Figure 9B). Statistical comparable *Tgfb* mRNA levels were found in WT CO and WT ONA (1.2-fold; p = 0.074) as well as in KO CO and KO ONA (0.7-fold, p = 0.07; Figure 9B). The evaluation of both immunized genotypes showed a significant reduction of *Tnfa* (0.5-fold, p = 0.036) and a significant increase of *Tgfb* (1.2-fold, p = 0.005) after immunization in KO mice (Figure 9C).

**Figure 9:**
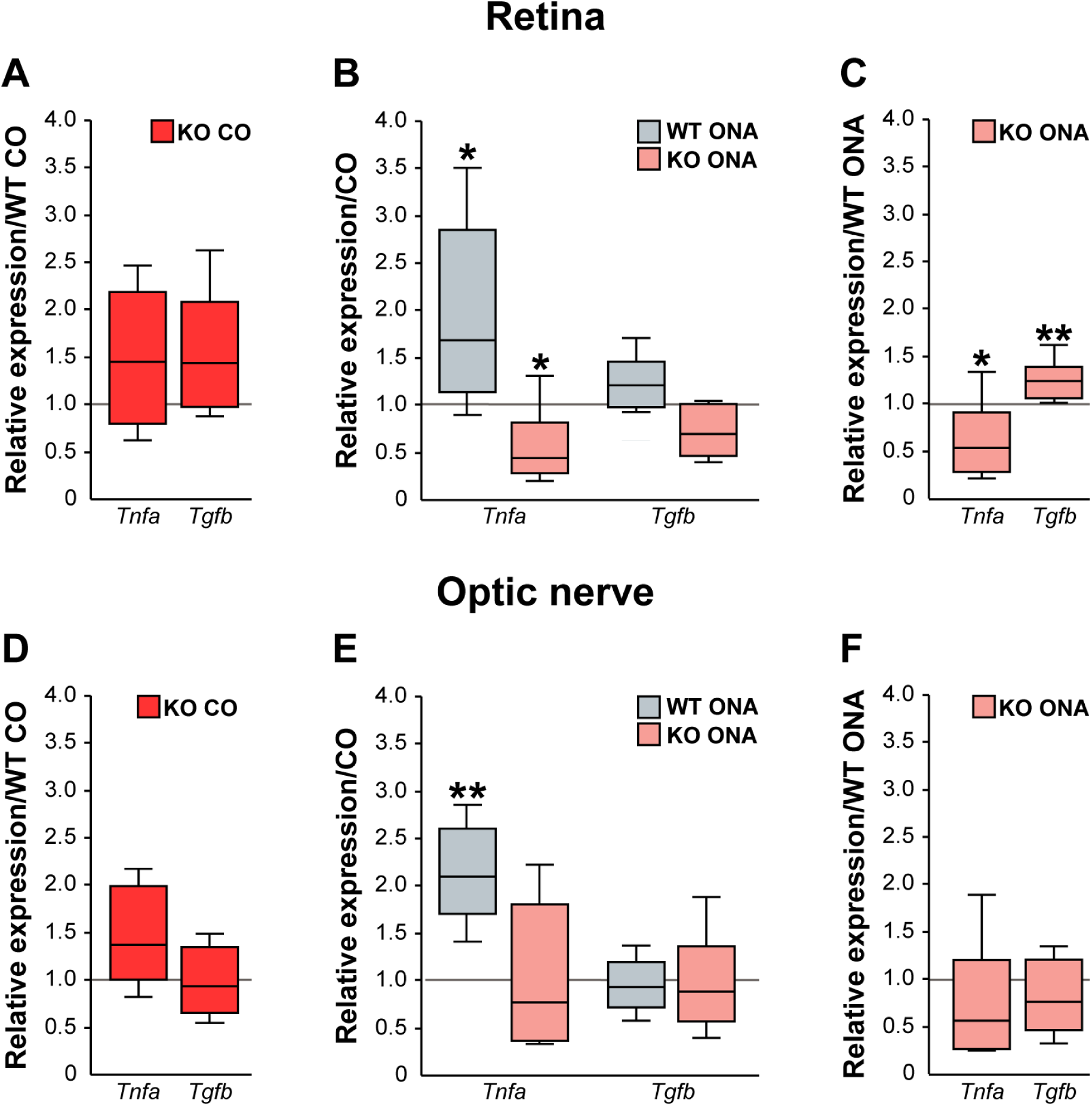
Reduced pro-inflammatory and enhanced anti-inflammatory cytokine expression in immunized KO mice. Relative expression of pro-inflammatory *Tnfa* and anti-inflammatory cytokine *Tgfb* was examined via RT-qPCR in control and immunized WT and KO retinae **(A-C)** and optic nerve tissue **(D-F). (A)** Analysis revealed comparable levels of *Tnfa* and *Tgfb* in KO CO compared to WT CO. **(B)** *Tnfa* expression level was significantly increased in WT ONA compared to WT CO. Strikingly, a reduced *Tnfa* mRNA level was found in KO ONA compared to the corresponding control group. Regarding *Tgfb*, the expression was comparable in both genotypes. **(C)** Comparison of WT ONA und KO ONA. Here, the pro-inflammatory factor was downregulated, and the anti-inflammatory cytokine was significantly upregulated in KO mice after immunization. **(D-F)** Similar expression patterns of both examined cytokines in optic nerve tissue. A significantly enhanced *Tnfa* expression could be detected in WT ONA compared to WT CO **(E)**. Values were shown as median ± quartile ± maximum/minimum. *p < 0.05; **p < 0.01. n = 5/group.

Finally, we examined, which microglial subtypes are altered due to an increased microglial reactivity in the optic nerve tissue of immunized and non-immunized WT and KO mice (Figure 9D-F, Supplementary Table 4). Equal mRNA levels of *Tnfa* (1.4-fold, p = 0.07) and *Tgfb* (0.9-fold, p = 0.659) were seen in WT and KO controls (Figure 9D). A significant enhanced *Tnfa* expression (2.1-fold, p = 0.008) and an unchanged *Tgfb* expression (0.9-fold, p = 0.71) were detected in WT ONA compared to WT CO (Figure 9E). No changes of these markers were found in KO CO in comparison to KO ONA mice (*Tnfa*: 0.8-fold, p = 0.443 and *Tgfb*: 0.9-fold, p = 0.575). In line with the RT-qPCR results of retinal tissue, we found a slightly reduced *Tnfa* (0.6-fold, p = 0.101) and an unaltered *Tgfb* (0.8-fold, p = 0.297) expression between both immunized groups (Figure 9F).

In conclusion, missing Tnc resulted in a reduced mRNA level of pro-inflammatory *Tnfa*, but an enhanced expression of anti-inflammatory *Tgfb*. The increased expression of the pro-inflammatory cytokine in WT after immunization points towards an enhanced presence of reactive M1 microglia.

## 4 Discussion

Glaucoma involves progressive degeneration of RGCs and their axons leading to visual field loss (Tochel et al., 2005; Shon et al., 2014; McMonnies, 2017). Developing glaucoma is often associated with elevated IOP, but RGC damage can also occur without IOP changes. Previous studies showed evidence that an altered immune response is involved in glaucoma pathology (Wax et al., 1998; Tezel et al., 1999; Wax et al., 2001; Joachim et al., 2003; Grus et al., 2006). In addition, remodeling of ECM constituents was found in several retinal neurodegenerative diseases, including glaucoma (Hernandez et al., 1990; Hernandez, 1992; Pena et al., 1999; Johnson et al., 2007; Reinhard et al., 2017a).

In the glaucoma pathology the mechanisms are poorly understood, especially the relationship between RGC loss and the role of the immune system as well as ECM molecules. Based on these findings, we characterized glaucomatous damage associated with the absence of the ECM glycoprotein Tnc in an EAG mouse model for the first time. Here, we immunized WT and KO mice with an optic nerve homogenate to induce retinal damage and analyzed IOP, retinal functionality, RGC degeneration, glial activation and pro- and anti-inflammatory cytokine expression 10 weeks later.

Our analyses revealed that the IOP of WT and KO stayed in normal ranges. Previous studies of the EAG animal model also showed no alteration in the IOP (Noristani et al., 2016; Reinehr et al., 2019). Comparable IOP in control and immunized animals, points to the fact that the EAG model can be a suitable model for normal tension glaucoma.

Analyses of retinal functionality via ERG recordings showed no differences in a- and b-wave responses in control and immunized WT and KO mice, which indicates that outer photoreceptor cells as well as bipolar/Müller glia cells are not affected in the EAG model.

Next, it should be investigated whether there is a glaucoma-typical damage, namely RGC loss in WT and KO animals after ONA-immunization. Our results demonstrated a significant loss of Brn3a^+^ RGCs in both genotypes following immunization. Interestingly, immunized KO mice displayed approximately 15 % more RGCs in retinal flat-mounts compared to immunized WT mice. Moreover, a comparable number of RGCs was found in retinal cross-sections of KO ONA. These findings indicate that Tnc deficiency leads to protection of RGCs in the EAG model. This is in line with our findings regarding optic nerve degeneration. Here, we verified a severe optic nerve damage by diminished βIII tubulin staining in immunized WT, while no alterations were found in KO ONA. Glaucomatous neurodegeneration was also evident in a pilot study of the EAG mouse model. In this study, a decrease of RGCs and degeneration of optic nerve was detected 6 weeks after immunization with different ONA-doses (Reinehr et al., 2019).

In the EAG rat model an early upregulation of Tnc and its interaction partner RPTPβ/ζ/phosphacan was observed at 7 days, whereas a significant loss of RGCs could be detected 22 days after immunization (Joachim et al., 2014; Reinehr et al., 2016). These results suggest that Tnc may serve as an early indicator of retinal damage.

Studies reported that Tnc is a pro-inflammatory mediator, which is involved in the pathogenesis of CNS autoimmunity (Jakovcevski et al., 2013; Momcilovic et al., 2017; Wiemann et al., 2019). In addition, an enhanced reactivity of astrocytes in response to inflammatory signals is directly regulated by the ECM (Johnson et al., 2015). Furthermore, degeneration of RGCs is accompanied by reactive astrogliosis, which results in a higher GFAP expression (Senatorov et al., 2006; Inman and Horner, 2007; Johnson and Morrison, 2009; Noristani et al., 2016; Reinhard et al., 2019). Here, we detected an enhanced GFAP level in WT mice after immunization. However, the lack of Tnc resulted in the loss of a significant astrocytic response, which might depend on a missing Tnc mediated pro-inflammatory signaling.

In this study we noted a reduced population of mature oligodendrocytes and OPCs after immunization in WT mice. Also, a reduced MBP immunoreactivity demonstrated a demyelination of the optic nerve in the WT condition. Furthermore, no evidence of a decrease in oligodendroglia density and demyelination was detected after ONA-immunization in KO animals.

Interestingly, the fraction of mature Olig2/CC1 double immunopositive oligodendrocytes was significantly increased in the KO, in agreement with our earlier finding that Tnc inhibits the maturation of oligodendrocytes (Czopka et al., 2009; Czopka et al., 2010). In addition, a strong expression of Tnc in the optic nerve head inhibits the migration of oligodendrocytes from the optic nerve into the retina (Bartsch et al., 1994). Based on our results and the mentioned studies, we suggest that Tnc has an impact on demyelination processes. Missing of Tnc leads to no alteration in oligodendroglia but has a protective effect on myelination of optic nerve fibers in our autoimmune induced glaucoma mouse model.

Neuroinflammatory changes in the retina occur during glaucomatous damage (Williams et al., 2017). Microglial activation is a very early event, often before significant loss of RGCs takes place (Ebneter et al., 2010; Bosco et al., 2011; Ramirez et al., 2017). An enhanced microglial infiltration and marker expression revealed an increased glial response in WT post immunization in our study. Moreover, a significantly increased expression of the pro-inflammatory cytokine *Tnfa* implied a higher activity of M1-microglia. Previous studies showed that microglia switched to a M1-like phenotype, which can lead to neurotoxic effects by producing high levels of pro-inflammatory cytokines (Block et al., 2007; Varnum and Ikezu, 2012; Tang and Le, 2016). It is already known that Tnc supports the activity of M1-microglia (Claycomb et al., 2014; Wiemann et al., 2019). In primary microglia Tnc induced release of *Tnfa* and regulated the expression of *iNOS* via Toll-like receptor 4 signaling (Haage et al., 2019). In glaucoma, a glia-derived neuronal death was described through TNF-α (Tezel et al., 2001; Kitaoka et al., 2006; Nakazawa et al., 2006; Cueva Vargas et al., 2015). However, in an early phase after an optic nerve crush a protective effect of TNF-α was found in an experimental animal model (Mac Nair et al., 2014).

In contrast, we found less invasion und reactivity of microglia in immunized KO mice and a significantly enhanced mRNA level of the anti-inflammatory *Tgfb*. Furthermore, the lack of Tnc leads to a significantly decreased expression of *Tnfa*. This result is consistent with the study by Piccinini, which demonstrated that the deletion of Tnc impaired TNF-α production (Piccinini and Midwood, 2012). TGF-β inhibits pro-inflammatory cytokines and regulates proliferation and activity of microglial cells (Piras et al., 2012). Moreover, a missing TGF-β signaling in microglia promotes retinal degeneration (Ma et al., 2019). Taken together, in our study an immunization induced increased microglia and inflammatory cytokine release, which was attenuated by the lack of Tnc. This finding strongly indicates that Tnc might be involved in glaucoma degeneration by regulating microglia reactivity and cytokine secretion.

Collectively, the loss of Tnc resulted in a reduced RGC loss, diminished macro- and microglial responses, and a shift towards an enhanced anti-inflammatory at the expense of pro-inflammatory signaling. Our results showed that Tnc deficiency has multiple neuroprotective effects, suggesting that Tnc signaling plays an important role in glaucomatous neurodegeneration.

## 5 Conclusion

Our study demonstrated that Tnc influences glial response, migration, and reactivity during glaucomatous damage. This model is ideally suited for a better understanding of the molecular mechanisms between retinal neurodegeneration and ECM remodeling in order to develop future therapeutic options.

## 6 Abbreviations

Brn3a: brain-specific homeobox/POU domain protein 3a
CC1: coiled coil-1
Cd68: cluster of differentiation 68
EAG: experimental autoimmune glaucoma
ECM: extracellular matrix
ERG: electroretinogram
GCL: ganglion cell layer
GFAP: glial fibrillary acidic protein
Iba1: ionized calcium-binding adapter molecule 1
INL: inner nuclear layer
IOP: intraocular pressure
IPL: inner plexiform layer
KO: knockout
KO CO: control group tenascin-C knockout
KO ONA: immunized tenascin-C knockout
MBP: myelin basic protein
NFL: nerve fiber layer
Nos2: nitric oxide synthase 2
Olig2: oligodendrocyte transcription factor 2
ONA: optic nerve antigen
ONL: outer nuclear layer
OPC: oligodendrocytes precursor cell
OPL: outer plexiform layer
RGC: retinal ganglion cell
*Tgfb*/TGF-β: transforming growth factor-beta
Tnc: tenascin-C
*Tnfa*/TNF-α: tumor necrosis factor-alpha
WT: wild type
WT CO: control group wild type
WT ONA: immunized wild type

## 7 Acknowledgement

The authors thank Stephanie Chun, Anja Coenen, Sabine Kindermann, Franziska Mennes, Annalena Pamp and Marion Voelzkow for excellent technical assistance.

## 9 Authors contribution

SW performed experiments, analyzed data and wrote the manuscript. JR designed the study, analyzed data and revised the manuscript. SR and ZC performed experiments and analyzed data. SCJ and AF designed the study and revised the manuscript. All authors read and approved the final manuscript.

## 11 Contribution to the field statement

Glaucoma is a complex, multifactorial disease, where retinal ganglion cell loss, optic nerve degeneration, glial activation, neuroinflammation and extracellular matrix remodeling are linked to glaucomatous damage. Previous studies described an altered immune response in glaucoma pathology. In the present study, we used an intraocular pressure independent experimental autoimmune glaucoma model to analyze the effect of the extracellular matrix glycoprotein tenascin-C on retinal ganglion cells and glial activity in glaucoma. In wild type animals, we verified severe damage of retinal ganglion cells, demyelination and reactive astrogliosis. Autoimmune-induced glaucomatous damage was also associated to neuroinflammation, characterized by an enhanced microglia infiltration and expression of reactive glial markers. In contrast, the lack of tenascin-C resulted in a reduced retinal ganglion cells loss and ameliorated demyelination of the optic nerve. Remarkably, absence of tenascin-C led to a missing macro- and microglial response and anti-inflammatory cytokine expression. Collectively, our results indicate that tenascin-C plays a significant role in glial response and neuroinflammatory processes during glaucomatous degeneration. Thus, this transgenic experimental autoimmune glaucoma mouse model offers for the first time the possibility to investigate intraocular pressure independent glaucomatous damage in direct relation to extracellular matrix remodeling.

