## Supplementary table 1 for "Loss of the extracellular matrix molecule tenascin-C leads to absence of reactive gliosis and promotes anti-inflammatory cytokine expression in an autoimmune glaucoma mouse model"

### *Supplementary Material*

#### 1 Supplementary Tables

**Supplementary Table 1:**
IOP measurements before and after immunization in WT and KO mice.

| Genotype | Group | Age (weeks) | Mean | SEM | P-value | N |
|---|---|---|---|---|---|---|
| WT | - | 5 | 9.8 | 0.2 | 1.0 | 16 |
| KO | - | 5 | 9.7 | 0.1 | | |
| WT | CO | 6 | 9.4 | 0.2 | >0.05 | 8 |
| WT | ONA | 6 | 9.2 | 0.3 | | |
| KO | CO | 6 | 9.1 | 0.3 | | |
| KO | ONA | 6 | 10.3 | 0.3 | | |
| WT | CO | 7 | 9.8 | 0.4 | >0.05 | 8 |
| WT | ONA | 7 | 9.9 | 0.4 | | |
| KO | CO | 7 | 9.9 | 0.3 | | |
| KO | ONA | 7 | 9.9 | 0.2 | | |
| WT | CO | 8 | 11.2 | 0.5 | >0.05 | 8 |
| WT | ONA | 8 | 11.3 | 0.3 | | |
| KO | CO | 8 | 10.1 | 0.3 | | |
| KO | ONA | 8 | 10.2 | 0.4 | | |
| WT | CO | 9 | 10.2 | 0.3 | >0.05 | 8 |
| WT | ONA | 9 | 10.9 | 0.4 | | |
| KO | CO | 9 | 10.8 | 0.3 | | |
| KO | ONA | 9 | 10.2 | 0.6 | | |
| WT | CO | 10 | 10.1 | 0.4 | >0.05 | 8 |
| WT | ONA | 10 | 9.7 | 0.3 | | |
| KO | CO | 10 | 10.6 | 0.4 | | |
| KO | ONA | 10 | 10.5 | 0.4 | | |
| WT | CO | 11 | 11.0 | 0.3 | >0.05 | 8 |
| WT | ONA | 11 | 10.6 | 0.2 | | |
| KO | CO | 11 | 10.9 | 0.7 | | |
| KO | ONA | 11 | 10.6 | 0.3 | | |
| WT | CO | 12 | 10.8 | 0.2 | >0.05 | 8 |
| WT | ONA | 12 | 10.8 | 0.3 | | |
| KO | CO | 12 | 10.3 | 0.4 | | |
| KO | ONA | 12 | 10.2 | 0.3 | | |
| WT | CO | 13 | 10.2 | 0.3 | >0.05 | 8 |
| WT | ONA | 13 | 9.5 | 0.3 | | |
| KO | CO | 13 | 10.5 | 0.4 | | |
| KO | ONA | 13 | 10.3 | 0.4 | | |
| WT | CO | 14 | 9.4 | 0.1 | >0.05 | 8 |
| WT | ONA | 14 | 9.4 | 0.2 | | |

| | | | | | | |
|---|---|---|---|---|---|---|
| KO | CO | 14 | 9.3 | 0.3 | | |
| KO | ONA | 14 | 10.8 | 0.4 | | |
| WT | CO | 15 | 9.7 | 0.2 | >0.05 | 8 |
| WT | ONA | 15 | 10.3 | 0.2 | | |
| KO | CO | 15 | 10.1 | 0.5 | | |
| KO | ONA | 15 | 10.2 | 0.5 | | |

**Supplementary Table 2:**
Data of a- and b-wave amplitudes recorded from WT CO, WT ONA, KO CO, and KO ONA animals. Values of light flash intensity (cd x s/m²) are displayed as mean ± SEM (n = 5/group).

| **Light flash intensity [cd x s/m²]** | **0.1** | | **0.3** | | **1** | | **3** | | **10** | | **25** | |
|---|---|---|---|---|---|---|---|---|---|---|---|---|
| **Amplitude [µV]** | **Mean** | **SEM** | **Mean** | **SEM** | **Mean** | **SEM** | **Mean** | **SEM** | **Mean** | **SEM** | **Mean** | **SEM** |
| **A-wave** | | | | | | | | | | | | |
| WT CO | 40.1 | 9.9 | 69.4 | 14.5 | 99.71 | 13.9 | 112.3 | 20.6 | 129.6 | 26.9 | 148.0 | 6.3 |
| WT ONA | 39.4 | 4.1 | 60.1 | 4.9 | 91.02 | 3.8 | 90.8 | 5.8 | 108.1 | 9.1 | 106.6 | 8.2 |
| KO CO | 27.4 | 3.2 | 41.4 | 7.2 | 68.87 | 13.9 | 87.6 | 13.6 | 114.5 | 20.1 | 103.8 | 26.5 |
| KO ONA | 35.8 | 2.6 | 67.8 | 7.2 | 80.80 | 4.7 | 92.3 | 3.3 | 109.9 | 14.8 | 151.6 | 16.2 |
| P-value | > 0.05 | | > 0.05 | | > 0.05 | | > 0.05 | | > 0.05 | | > 0.05 | |
| **B-wave** | | | | | | | | | | | | |
| WT CO | 279.4 | 32.9 | 335.7 | 43.3 | 333.98 | 41.5 | 302.2 | 41.9 | 355.1 | 48.2 | 354.8 | 37.9 |
| WT ONA | 250.6 | 21.9 | 331.3 | 27.5 | 348.31 | 32.4 | 365.5 | 41.1 | 356.4 | 35.7 | 381.9 | 48.9 |
| KO CO | 169.0 | 25.9 | 188.9 | 39.9 | 228.82 | 39.1 | 247.0 | 38.7 | 249.3 | 49.2 | 330.9 | 38.8 |
| KO ONA | 233.1 | 18.4 | 284.0 | 28.8 | 309.79 | 33.0 | 307.6 | 24.6 | 307.8 | 30.6 | 318.0 | 34.7 |
| P-value | > 0.05 | | > 0.05 | | > 0.05 | | > 0.05 | | > 0.05 | | > 0.05 | |

**Supplementary Table 3:**
Cell counts of Brn3a$^{+}$ and Iba1$^{+}$ cells [%] in WT CO, WT ONA, KO CO, and KO ONA animals. WT CO group was set to 100%. P-values < 0.05 are shown in bold.

| Geno-type | Group | Retinal area | Tissue | Mean | SEM | P-value | N |
|---|---|---|---|---|---|---|---|
| **Brn3a$^{+}$ cells [%]** | | | | | | | |
| WT | CO | - | Cross-section | 100.0 | 4.2 | **0.004** (WT CO vs. WT ONA)<br>0.64 (WT CO vs. KO CO)<br>0.10 (WT CO vs. KO ONA)<br>**0.04** (WT ONA vs. KO CO)<br>0.39 (WT ONA vs. KO ONA)<br>0.57 (KO CO vs. KO ONA) | 5 |
| WT | ONA | | | 73.1 | 6.1 | | |
| KO | CO | | | 92.2 | 3.9 | | |
| KO | ONA | | | 83.7 | 8.7 | | |
| WT | CO | Central | Flat-mount | 100.0 | 2.5 | **< 0.001** (WT CO vs. WT ONA)<br>0.94 (WT CO vs. KO CO)<br>**0.007** (WT CO vs. KO ONA) | 9 |
| | | Peripheral | | 100.0 | 1.7 | **< 0.001** (WT CO vs. WT ONA)<br>0.99 (WT CO vs. KO CO)<br>**0.01** (WT CO vs. KO ONA) | |
| | | Total | | 100.0 | 2.0 | **< 0.001** (WT CO vs. WT ONA)<br>0.96 (WT CO vs. KO CO)<br>**0.01** (WT CO vs. KO ONA) | |
| WT | ONA | Central | | 82.7 | 1.7 | **< 0.001** (WT ONA vs. WT CO)<br>**0.003** (WT ONA vs. KO CO)<br>0.80 (WT ONA vs. KO ONA) | 9 |
| | | Peripheral | | 77.0 | 1.8 | **< 0.001** (WT ONA vs. WT CO)<br>**< 0.001** (WT ONA vs. KO CO)<br>0.06 (WT ONA vs. KO ONA) | |
| | | Total | | 80.3 | 1.5 | **< 0.001** (WT ONA vs. WT CO)<br>**< 0.001** (WT ONA vs. KO CO)<br>0.27 (WT ONA vs. KO ONA) | |
| KO | CO | Central | | 97.8 | 2.9 | 0.94 (KO CO vs. WT CO)<br>**0.003** (KO CO vs. WT ONA)<br>**0.03** (KO CO vs. KO ONA) | 9 |
| | | Peripheral | | 99.0 | 4.1 | 0.99 (KO CO vs. WT CO)<br>**< 0.001** (KO CO vs. WT ONA)<br>**0.02** (KO CO vs. KO ONA) | |
| | | Total | | 98.2 | 3.3 | 0.96 (KO CO vs. WT CO)<br>**< 0.001** (KO CO vs. WT ONA)<br>**0.02** (KO CO vs. KO ONA) | |
| KO | ONA | Central | | 86.3 | 3.7 | **0.007** (KO ONA vs. WT CO)<br>0.80 (KO ONA vs. WT ONA)<br>**0.03** (KO ONA vs. KO CO) | 9 |
| | | Peripheral | | 87.1 | 2.8 | **0.01** (KO ONA vs. WT CO)<br>0.06 (KO ONA vs. WT ONA)<br>**0.02** (KO ONA vs. KO CO) | |
| | | Total | | 86.9 | 3.1 | **0.01** (KO ONA vs. WT CO) | |

| | | | | | | | |
|---|---|---|---|---|---|---|---|
| | | | | | | 0.27$^{\text{KO ONA vs. WT ONA}}$<br>**0.02**$^{\textbf{KO ONA vs. KO CO}}$ | |
| **Iba1$^{+}$ cells [%]** | | | | | | | |
| WT | CO | Central | Flat-mount | 100.0 | 3.5 | **0.002**$^{\textbf{WT CO vs. WT ONA}}$<br>0.99$^{\text{WT CO vs. KO CO}}$<br>**0.04**$^{\textbf{WT CO vs. KO ONA}}$ | 9 |
| | | Peripheral | | 100.0 | 3.2 | **0.002**$^{\textbf{WT CO vs. WT ONA}}$<br>0.62$^{\text{WT CO vs. KO CO}}$<br>0.08$^{\text{WT CO vs. KO ONA}}$ | |
| | | Total | | 100.0 | 2.9 | **< 0.001**$^{\textbf{WT CO vs. WT ONA}}$<br>0.97$^{\text{WT CO vs. KO CO}}$<br>**0.03**$^{\textbf{WT CO vs. KO ONA}}$ | |
| WT | ONA | Central | | 122.5 | 2.9 | **0.002** $^{\textbf{WT ONA vs. WT CO}}$<br>**< 0.001**$^{\textbf{WT ONA vs. KO CO}}$<br>**< 0.001**$^{\textbf{WT ONA vs. KO ONA}}$ | 9 |
| | | Peripheral | | 123.6 | 3.4 | **0.002** $^{\textbf{WT ONA vs. WT CO}}$<br>**< 0.05**$^{\textbf{WT ONA vs. KO CO}}$<br>**< 0.001**$^{\textbf{WT ONA vs. KO ONA}}$ | |
| | | Total | | 123.0 | 2.4 | **< 0.001**$^{\textbf{WT ONA vs. WT CO}}$<br>**0.002**$^{\textbf{WT ONA vs. KO CO}}$<br>**< 0.001**$^{\textbf{WT ONA vs. KO ONA}}$ | |
| KO | CO | Central | | 98.1 | 5.8 | 0.08$^{\text{KO CO vs. KO ONA}}$<br>0.99$^{\text{KO CO vs. WT CO}}$<br>**< 0.001**$^{\textbf{KO CO vs. WT ONA}}$ | 9 |
| | | Peripheral | | 107.2 | 6.4 | **0.004**$^{\textbf{KO CO vs. KO ONA}}$<br>0.63$^{\text{KO CO vs. WT CO}}$<br>**< 0.05**$^{\textbf{KO CO vs. WT ONA}}$ | |
| | | Total | | 102.3 | 5.7 | **0.009**$^{\textbf{KO CO vs. KO ONA}}$<br>0.97$^{\text{KO CO vs. WT CO}}$<br>**0.002**$^{\textbf{KO CO vs. WT ONA}}$ | |
| KO | ONA | Central | | 84.0 | 2.9 | 0.08$^{\text{KO ONA vs. KO CO}}$<br>**0.04**$^{\textbf{KO ONA vs. WT CO}}$<br>**< 0.001**$^{\textbf{KO ONA vs. WT ONA}}$ | 9 |
| | | Peripheral | | 85.1 | 3.1 | **0.004**$^{\textbf{KO ONA vs. KO CO}}$<br>0.08$^{\text{KO ONA vs. WT CO}}$<br>**< 0.001**$^{\textbf{KO ONA vs. WT ONA}}$ | |
| | | Total | | 84.5 | 2.7 | **0.009**$^{\textbf{KO ONA vs. KO CO}}$<br>**0.03**$^{\textbf{KO ONA vs. WT CO}}$<br>**< 0.001**$^{\textbf{KO ONA vs. WT ONA}}$ | |

**Supplementary Table 4:**
RTq-PCR analyses of glial cell types and pro- and anti-inflammatory cytokines in WT CO, WT ONA, KO CO, and KO ONA animals, the fold change of the expression is displayed. P-values < 0.05 are shown in bold (n = 5/group).

| Genotype/ group | Gene | Tissue | Median | Quartile + maximum/minimum | P-value |
|---|---|---|---|---|---|
| WT CO vs. KO CO | *Gfap* | Retina | 1.4 | 1.012 - 2.016 | 0.110 |
| | | | | 0.724 - 2.500 | |
| | | Optic nerve | 1.1 | 0.799 - 1.406 | 0.539 |
| | | | | 0.645 - 1.531 | |
| WT CO vs. WT ONA | | Retina | 1.7 | 1.011 - 2.751 | **0.044** |
| | | | | 0.902 - 3.380 | |
| | | Optic nerve | 1.4 | 0.970 - 1.923 | 0.071 |
| | | | | 0.845 - 2.083 | |
| KO CO vs. KO ONA | | Retina | 1.2 | 0.812 - 1.817 | 0.362 |
| | | | | 0.614 - 2.557 | |
| | | Optic nerve | 0.5 | 0.300 - 0.918 | **0.047** |
| | | | | 0.174 - 1.491 | |
| WT ONA vs. KO ONA | | Retina | 1 | 0.625 - 1.518 | 0.993 |
| | | | | 0.450 - 2.136 | |
| | | Optic nerve | 0.4 | 0.242 - 0.705 | **0.021** |
| | | | | 0.128 - 1.150 | |
| WT CO vs. KO CO | *Iba1* | Retina | 1.3 | 0.924 - 1.869 | 0.201 |
| | | | | 0.665 - 2.254 | |
| | | Optic nerve | 1.1 | 0.757 - 1.502 | 0.607 |
| | | | | 0.614 - 1.827 | |
| WT CO vs. WT ONA | | Retina | 1.5 | 1.124 - 2.100 | **0.048** |
| | | | | 0.816 - 2.941 | |
| | | Optic nerve | 1.5 | 1.117 - 2.068 | **0.032** |
| | | | | 0.860 - 2.515 | |
| KO CO vs. KO ONA | | Retina | 0.9 | 0.672 - 1.182 | 0.399 |
| | | | | 0.611 - 1.424 | |
| | | Optic nerve | 0.9 | 0.584 - 1.526 | 0.842 |
| | | | | 0.437 - 2.189 | |
| WT ONA vs. KO ONA | | Retina | 0.8 | 0.629 - 1.063 | 0.2 |
| | | | | 0.478 - 1.363 | |
| | | Optic nerve | 0.7 | 0.463 - 1.160 | 0.148 |
| | | | | 0.327 - 1.436 | |
| WT CO vs. KO CO | *Nos2* | Retina | 1.7 | 1.240 - 2.151 | **0.013** |
| | | | | 1.006 - 2.342 | |
| | | Optic nerve | 1.2 | 0.881 - 1.618 | 0.292 |
| | | | | 0.648 - 2.009 | |
| WT CO vs. WT ONA | | Retina | 1.4 | 1.144 - 1.892 | **0.021** |
| | | | | 0.873 - 2.408 | |
| | | Optic nerve | 1.4 | 1.095 - 1.633 | **0.008** |

| | | | | | |
|---|---|---|---|---|---|
| | | | | 0.975 - 1.888 | |
| KO CO vs. KO ONA | | Retina | 0.9 | 0.716 - 1.172<br>0.614 - 1.261 | 0.605 |
| | | Optic nerve | 0.6 | 0.372 - 1.224<br>0.238 - 1.829 | 0.197 |
| WT ONA vs. KO ONA | | Retina | 1.1 | 0.743 - 1.381<br>0.618 - 1.897 | 0.65 |
| | | Optic nerve | 0.6 | 0.298 - 1.093<br>0.236 - 1.377 | **0.043** |
| WT CO vs. KO CO | *Cd68* | Retina | 1.3 | 1.105 - 1.731<br>0.986 - 1.985 | **0.017** |
| | | Optic nerve | 1.2 | 0.956 - 1.644<br>0.758 - 2.009 | 0.149 |
| WT CO vs. WT ONA | | Retina | 1.3 | 1.101 - 1.732<br>0.893 - 1.976 | **0.032** |
| | | Optic nerve | 1.1 | 0.678 - 1.926<br>0.530 - 2.763 | 0.752 |
| KO CO vs. KO ONA | | Retina | 0.9 | 0.735 - 1.049<br>0.654 - 1.226 | 0.153 |
| | | Optic nerve | 0.9 | 0.572 - 1.504<br>0.444 - 2.403 | 0.685 |
| WT ONA vs. KO ONA | | Retina | 0.9 | 0.731 - 1.133<br>0.646 - 1.283 | 0.3 |
| | | Optic nerve | 1.0 | 0.534 - 1.846<br>0.323 - 3.403 | 0.93 |
| WT CO vs. KO CO | *Tnfa* | Retina | 1.4 | 0.795 – 2.186<br>0.612 – 2.466 | 0.132 |
| | | Optic nerve | 1.4 | 1.011 - 1.984<br>0.818 - 2.177 | 0.07 |
| WT CO vs. WT ONA | | Retina | 1.7 | 1.131 - 2.836<br>0.884 - 3.491 | **0.026** |
| | | Optic nerve | 2.1 | 1.712 - 2.615<br>1.414 - 2.868 | **0.008** |
| KO CO vs. KO ONA | | Retina | 0.4 | 0.276 - 0.810<br>0.187 - 1.305 | **0.031** |
| | | Optic nerve | 0.8 | 0.366 - 1.813<br>0.335 - 2.231 | 0.443 |
| WT ONA vs. KO ONA | | Retina | 0.5 | 0.288 - 0.916<br>0.210 - 1.335 | **0.036** |
| | | Optic nerve | 0.6 | 0.278 - 1.202<br>0.255 - 1.902 | 0.101 |
| WT CO vs. KO CO | *Tgfb* | Retina | 1.4 | 0.969 – 2.078<br>0.866 - 2.625 | 0.071 |
| | | Optic nerve | 0.9 | 0.660 - 1.349<br>0.554 - 1.491 | 0.659 |
| WT CO vs. WT | | Retina | 1.2 | 0.977 - 1.459 | 0.074 |

| | | | | | |
|---|---|---|---|---|---|
| ONA | | | | 0.923 - 1.707 | |
| | | Optic nerve | 0.9 | 0.716 - 1.204 | 0.71 |
| | | | | 0.583 – 1.372 | |
| KO CO vs. KO ONA | | Retina | 0.7 | 0.471 - 1.017 | 0.07 |
| | | | | 0.397 - 1.043 | |
| | | Optic nerve | 0.9 | 0.575 - 1.373 | 0.575 |
| | | | | 0.403 - 1.878 | |
| WT ONA vs. KO ONA | | Retina | 1.2 | 1.058 - 1.386 | **0.005** |
| | | | | 1.008 - 1.612 | |
| | | Optic nerve | 0.8 | 0.479 - 1.222 | 0.297 |
| | | | | 0.333 - 1.785 | |

#### 1.1 Supplementary Figures

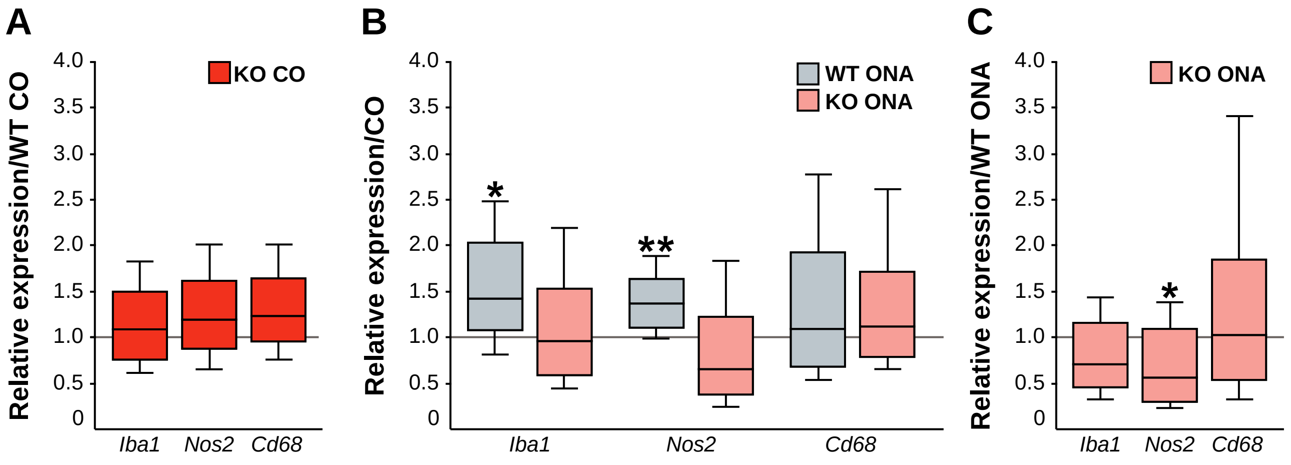


**Supplementary Figure 1:** RTq-PCR analyses of optic nerve tissue from control and immunized WT and KO mice.

**(A)** Examination of relative *Iba1*, *Nos2*, and *Cd68* mRNA expression showed no changes in KO CO compared to WT CO. **(B)** Compared to WT CO, a significant upregulation of *Iba1* and *Nos2* levels was verified in WT ONA. While no significant changes were detected regarding the expression levels of these markers in KO ONA compared to KO CO. **(C)** After immunization, a significantly downregulation of *Nos2* expression was observed in KO ONA compared to WT ONA, whereas comparable mRNA level of *Iba1 and Cd68* were detected in KO ONA vs. WT ONA. Groups were compared using the pairwise fixed reallocation and randomization test and were shown as median ± quartile ± minimum/maximum. n = 5/group. *p < 0.05; **p < 0.01.
